# Acute IL-6 exposure triggers canonical IL-6R signalling in hiPSC microglia, but not neural progenitor cells

**DOI:** 10.1101/2022.08.05.502958

**Authors:** Amalie C. M. Couch, Shiden Solomon, Alessia Marrocu, Rodrigo Duarte, Yiqing Sun, Laura Sichlinger, Rugile Matuleviciute, Lucia Dutan Polit, Bjørn Hanger, Shahram Kordasti, Deepak P. Srivastava, Anthony C. Vernon

**Author notes:** These authors share senior authorship (Deepak P. Srivastava, Anthony C. Vernon). **Corresponding author:** Dr. Anthony Vernon, Department of Basic and Clinical Neuroscience, Maurice Wohl Clinical Neuroscience Institute, Institute of Psychiatry, Psychology and Neuroscience, King’s College London, London SE5 9RT, United Kingdom.

## Abstract

**Background:** Exposure to elevated interleukin (IL)-6 levels *in utero* is consistently associated with increased risk for psychiatric disorders with a putative neurodevelopmental origin, such as schizophrenia (SZ) and autism spectrum condition (ASC). Although rodent models provide causal evidence for this association, we lack a detailed understanding of the cellular and molecular mechanisms in human model systems. To close this gap, we characterised the response of hiPSC-derived microglia-like cells (MGL) and neural progenitor cells (NPCs) to IL-6 in monoculture.

**Results:** We observed that human forebrain NPCs did not respond to acute IL-6 exposure in monoculture at both a protein and transcript level due to the absence of *IL-6Ra* expression and sIL-6Ra secretion. By contrast, acute IL-6 exposure resulted in STAT3 phosphorylation and increased *IL-6, JMJD3* and *IL-10* expression in MGL, confirming activation of canonical IL-6R signalling. Bulk RNAseq identified 156 upregulated genes (FDR <0.05) in MGL following acute IL-6 exposure, including *IRF8, REL, HSPA1A/B* and *OXTR*, which significantly overlapped with an upregulated gene set from *post-mortem* brain tissue from individuals with schizophrenia. Acute IL-6 stimulation significantly increased MGL motility suggestive of a gain of surveillance function, consistent with gene ontology pathways highlighted from the RNAseq data. Finally, MGLs displayed elevated CCL1, CXCL1, MIP-1A/B, IL-8, IL-13, IL-16, IL-18, MIF and Serpin-E1 secretion post 3h and 24h IL-6 exposure.

**Conclusion:** Our data provide evidence for cell specific effects of acute IL-6 exposure in a human model system and strongly suggest microglia-NPC co-culture models are required to study how IL-6 influences human cortical neural progenitor cell development *in vitro*.

## Introduction

Maternal immune activation (MIA) during pregnancy is consistently associated with an increased risk of neurodevelopmental disorders (NDDs) in the offspring, including schizophrenia (SZ) and autism spectrum condition (ASC) (Estes and McAllister, 2016). Evidence from both human and rodent studies suggests that the maternal peripheral cytokine profile mediates, at least in part, this increased risk for NDD-relevant phenotypes in the offspring (Allswede et al., 2020; Careaga et al., 2017; Graham et al., 2018; Meyer, 2014; Mueller et al., 2021; Rasmussen et al., 2021, 2019; Rudolph et al., 2018). Of this cytokine milieu, data from human, rodents and *in vitro* models have consistently identified interleukin (IL-) 6 as a sensor, effector, and transducer of environmental risk factor exposures, including MIA, on the fetal brain (Graham et al., 2018; Ozaki et al., 2020; Perry et al., 2021; Rasmussen et al., 2021, 2019; Rudolph et al., 2018; Smith et al., 2007). Specifically, in Mendelian randomization (MR) studies, genetically predicted IL-6 is associated with increased risk for schizophrenia in univariable MR (Perry et al., 2021). Elevated IL-6 concentrations in maternal serum correlate with modified amygdala and frontolimbic white matter connectivity in the offspring, which influence both cognitive development and some externalizing behaviors in the offspring (Graham et al., 2018; Rasmussen et al., 2021, 2019; Rudolph et al., 2018). In a mouse model of MIA based on maternal exposure to the viral mimetic Poly I:C, transcripts of *IL-6* are shown to be consistently elevated in maternal liver, placenta, and fetal primary microglia (Ozaki et al., 2020). Furthermore, peripheral IL-6 levels remain elevated in mouse offspring identified as susceptible to MIA, based on behavioral deficits relevant for SZ and ASC (Mueller et al., 2021). Moreover, acute elevation of IL-6 by injection into pregnant mice or developing embryos enhances glutamatergic synapse development resulting in overall brain hyperconnectivity and behavioral deficits relevant for ASC in adult offspring (Mirabella et al., 2021). Finally, blocking IL-6 in the pregnant rodent dam, irrespective of the immune stimulation paradigm, eliminates the pathological effects of MIA in the fetal rodent brain and subsequent behavioral deficits in the adult animal (Smith et al., 2007).

Human epidemiological or neuroimaging studies cannot however establish the cellular or molecular basis of IL-6 effects. Whilst animal models address this gap, the extent to which data from such models may translate to humans remains unclear, due to the species-specific gene regulation networks that encompass human neurodevelopment (Yokoyama et al., 2014). This is compounded by heterogeneity between laboratories in the gestational timing, dose, frequency, and route of administration of the infectious challenge in rodents (Smolders et al., 2018) and batch-to-batch heterogeneity of infectious agents (Mueller et al., 2019). As such, conflicting findings exist in the animal MIA literature regarding cellular mechanisms, exemplified by studies on the role of microglia (Smolders et al., 2018). Third, only a tiny fraction of animal studies have investigated cellular or molecular phenotypes proximal to the MIA event in the developing brain. This is important as knowledge of the most proximal molecular events to MIA could reveal important therapeutic targets for prevention of downstream pathology.

Human induced pluripotent stem cells (hiPSC), which may be differentiated into multiple different neural and glial lineages, have the potential to address these gaps in our knowledge. The power of hiPSC as a tool for investigating immune-associated mechanisms that occur in early brain development linked to NDD is evidenced by existing studies. Specifically, hiPSC directed towards neuronal fates have been utilized to investigate the pathological impact of Zika virus infection (Muffat et al., 2018), exposure to TLR3-agonists (Ritchie et al., 2018), and following direct exposure to cytokines, including interferon-gamma (Warre-Cornish et al., 2020) and IL-6 (Kathuria et al., 2022a). These studies are however critically lacking in one key aspect in that they have exclusively focused on neurons or astrocytes, at the expense of human microglia (Bhat et al., 2022; Park et al., 2020; Russo et al., 2018; Warre-Cornish et al., 2020). We therefore lack data on the impact of IL-6 on these critical immune-effector cells in a human–relevant model. Converging lines of evidence from human genetics, brain *post-mortem* tissue studies, neuroimaging and peripheral biomarker studies implicate microglia and the innate immune system in the pathophysiology of NDDs (Coomey et al., 2020; Mondelli et al., 2017). Since microglia also play critical roles in shaping neurodevelopment and the central immune response to maintain homeostasis (Hanger et al., 2020; Paolicelli et al., 2011), incorporating human microglia into hiPSC models to study the effects of immune activation on development is vital (Gonzalez et al., 2017; Russo et al., 2018).

To this end, we evaluated whether, and how, hiPSC-derived microglia like cells (MGLs) and neural progenitor cells (NPCs) respond to acute IL-6 stimulation in monocultures. We considered the following four questions: (1) do the cells have the receptor machinery to respond to IL-6 and other cytokines; (2) do these cells respond to acute IL-6; (3) does acute IL-6 induce a transcriptional profile similar to that seen in NDDs; and finally, (4) how does acute IL-6 impact the function of human MGLs?

## Results

### Human iPSC-derived microglial-like cells express IL-6 signaling receptors, but cortical neural progenitor cells do not

Successful differentiation of hiPSC to MGLs and forebrain NPCs was confirmed by expression of key signature genes and proteins for each cell type (**Supplementary Figure 1**). We then profiled human iPSC derived MGL and NPC monocultures (N=3 neurotypical male donors with N=3 separate clones per donor), for cytokine receptor expression by qPCR to establish the potential of each cell type to respond to IL-6, and other cytokines, *in vitro*. Transcript expression of *IFNyR1/2, TNFARSF1A, IL- 17Ra*, and both subunits required for IL-6 signaling, *IL-6Ra* and *IL-6ST*, all significantly increased with longer differentiation of MGLs *in vitro* relative to the hiPSC state (**Figure 1A**, statistics in **Supplementary Table 11**). Expression of *TNFRSF1B* was not significantly different from the hiPSC stage overall but numerically increased throughout MGL differentiation up to day 14 (**Figure 1A**). These data indicate that MGLs would be responsive to at least IL-6, IFN_γ_, TNF_α_ and IL-17.

**Figure 1:**
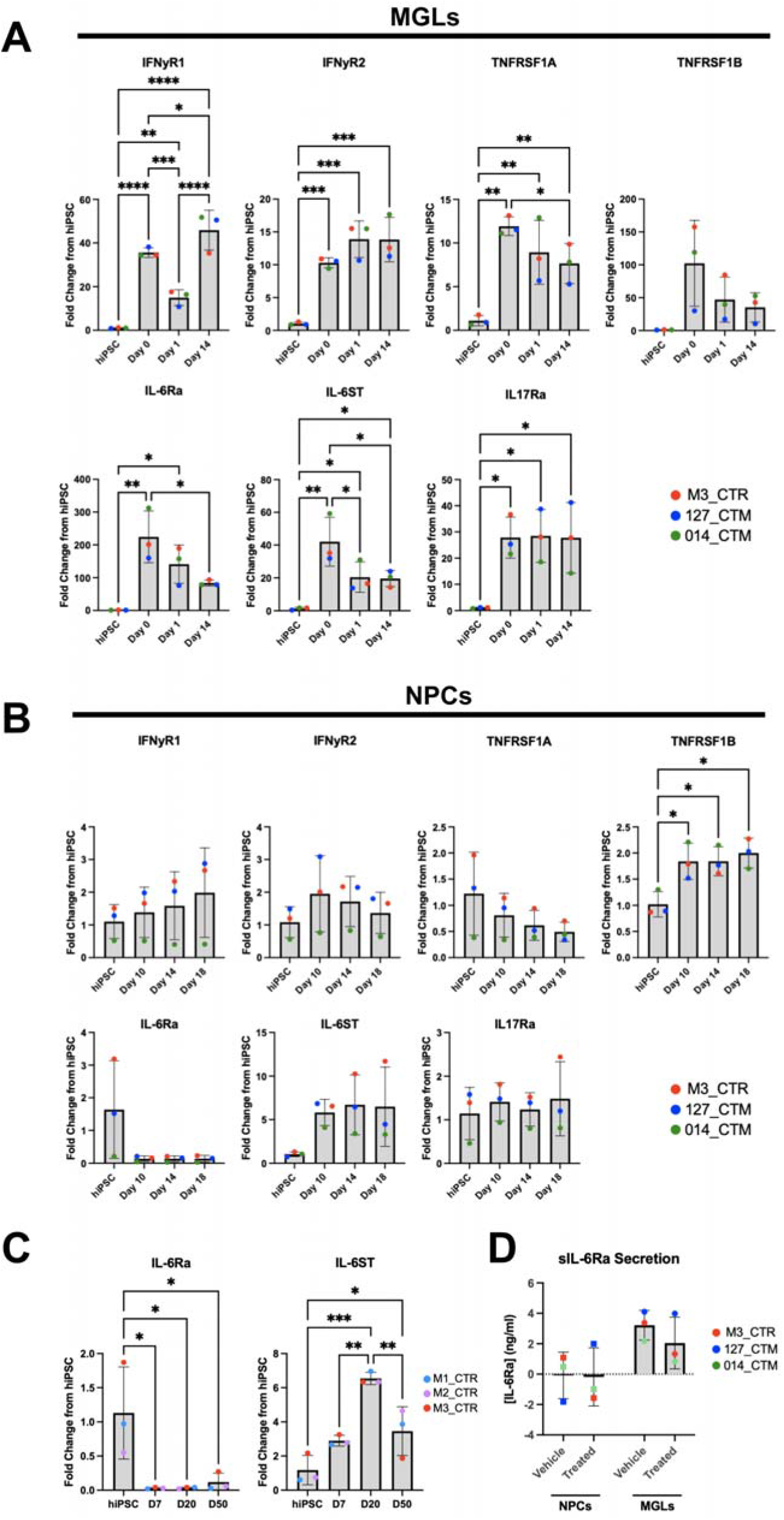
Cytokine receptor transcript expression in MGL and NPCs. (**A**) MGL differentiation time-course of cytokine receptors (*IFNyR1, IFNyR2, TNFRSF1A, TNFRSF1B, IL-6Ra, IL-6ST and IL- 17Ra*) by qPCR RNA samples at hiPSC, myeloid factory (day 0), MGL progenitor (day 1) and MGL (day 14) of differentiation. M3_CTR day 14 *IL17Ra* data point averaged from N=2 clones only. (**B**) NPC differentiation time-course of the same cytokine receptors by qPCR RNA samples at hiPSC and days 10, 14 and 18 of neuralisation to NPCs. 127_CTM day 14 *TNFRSF1B* data point was averaged form N=2 clones only. (**C**) qPCR of IL-6Ra and IL-6ST transcript expression in N=3 different healthy male lines (M1_CTR, M2_CTR and M3_CTR, one technical repeat each) over a longer timeframe, from hiPSC to D50 mature neurons. (**D**) Protein concentrations quantified by ELISA of soluble IL-6R (ng/ml) in NPC and MGL culture media after 3h vehicle or IL-6 100ng/ml exposure. Data shown are from N=3 male neurotypical hiPSC cell lines, averaged from three technical replicate clones per donor, unless stated otherwise where lines were outliers were excluded. 5% FDR BH method corrections after one-way ANOVA formatted as follows: *p < 0.05, **p < 0.01, ***p < 0.001, and ****p < 0.0001; not significant not labelled. Bar graphs plotted as mean with standard deviation (SD) error bars, and points coloured by donor line as shown in key.

In contrast to MGLs, differentiation of hiPSC to forebrain NPCs in monoculture led to virtually undetectable levels of *IL-6Ra* expression, relative to the hiPSC stage (**Figure 1B** statistics in **Supplementary Table 12**). This observed loss of *IL-6Ra* expression upon differentiation is in good agreement with previously reported data from human *post-mortem* brain tissue (Zhang et al., 2016). Specifically, publicly available gene expression datasets obtained at fetal and adult stages demonstrate *IL-6Ra* is primarily expressed by microglia and in part by astrocytes in the human brain, but not in neurons (Miller et al., 2014; Zhang et al., 2016) (**Supplementary Table 13**). Transcripts for *IFNyR1/2, TNFRF1A, IL-6ST* and *IL-17Ra* were expressed in forebrain NPCs, with expression levels remaining constant throughout neuralization, except for *TNFRSF1B*, which increased significantly relative to the hiPSC state (**Figure 1B**, statistics in **Supplementary Table 12**). These data indicate forebrain NPCs may be unresponsive to IL-6 when grown in monoculture, via *cis*- IL6R signaling, but responsive to IFNγ as shown previously (Warre-Cornish et al., 2020).

A recent study provides data to suggest that transcriptional and morphological phenotypes may be induced in hiPSC-derived mature cortical pyramidal neurons following exposure to IL-6 (Kathuria et al., 2022b). We therefore sought to replicate our finding of low to undetectable *IL-6Ra* expression and extend this analysis to longer differentiation times using an identical dual SMAD inhibition protocol, with different hiPSC lines from male, neurotypical donors (N=3). Analysis of RNA samples by qPCR confirmed the absence of *IL-6Ra* expression in forebrain NPCs and provides evidence to suggest this continues to be the case in mature neurons, at least at 50-days of differentiation *in vitro* (1-way ANOVA p=0.0112; **Figure 1C**). By contrast, *IL-6ST* expression increased throughout all stages of differentiation (1-way ANOVA p=0.0005; **Figure 1C**). These data suggest the absence of *IL-6Ra* expression in forebrain NPCs and mature neurons is neither donor line nor time point specific under the conditions tested.

Given the apparent loss of expression of *IL-6Ra* in forebrain NPCs at a transcript level, we next examined the secretion of the soluble IL-6Ra protein in both forebrain NPCs and MGLs after a 3h exposure to IL-6, indicative of the ability to initiate *trans*- IL-6 signaling (**Figure 1D**) (Campbell et al., 2014; Michalopoulou et al., 2004; Wolf et al., 2014). Using a custom-ELISA kit in vehicle-treated cultures, we observed no secretion of sIL-6Ra protein into the culture supernatant by forebrain NPCs, with over half of the measurements taken below the detectable range (**Figure 1D**). By contrast, sIL-6Ra secretion was clearly present in vehicle-treated MGLs in the culture supernatant (**Figure 1D**). Acute (3 hr) exposure to IL-6 (100 ng/ml) did not increase the secretion of sIL-6Ra protein however, from either cell type (2-Way ANOVA: cell type F=9.349; p=0.0156, treatment F=0.4969; p=0.5009, interaction F=0.3499; p=0.5709). Nonetheless, the fact that MGLs secrete sIL-6Ra provides an opportunity for other cell types within their vicinity to respond to IL-6 via *trans* signaling. By contrast, the lack of sIL-6Ra secretion from NPCs confirms the fact they do not express the soluble form of the IL-6 receptor and strongly suggests forebrain NPCs are unlikely to be responsive to IL-6 in *monoculture* under the conditions tested.

### Microglial-like cells, but not forebrain NPCs activate the canonical STAT3 pathway after IL-6 exposure in monoculture

We next determined the functionality of the IL-6Ra and IL-6ST receptors in MGL progenitors, MGLs and NPCs (**Figure 2A**). First, transcripts of relevant STAT3 downstream target genes (*IL-6, TNF*_α_, *JMJD3* and *IL-10*) (Przanowski et al., 2014) were measured by qPCR (**Figure 2B**, statistics in **Supplementary Table 14**). At 3h after exposure to IL-6 (100 ng/ml), both MGL progenitor and MGL cells responded to IL-6 by increasing the expression of *IL-6* itself and *JMJD3* transcript expression, relative to the vehicle control. In addition, IL-6 exposed MGLs, but not MGL progenitors, increased the expression of *IL-10*, whilst expression of *TNF*_α_ was not affected in either MGL progenitors or MGLs (**Figure 2B**, statistics in **Supplementary Table 14**). By contrast, at 24h after exposure to IL-6 (100 ng/ml), the expression of all these genes was no longer significantly different relative to the vehicle control in both MGL progenitors and MGLs (**Figure 2B**). Based on the apparent functional maturity of MGLs after 14 days of differentiation (Haenseler et al., 2017), and their response to IL-6 by increasing *IL-10*, only MGLs differentiated for 14 days were used in all subsequent experiments.

**Figure 2:**
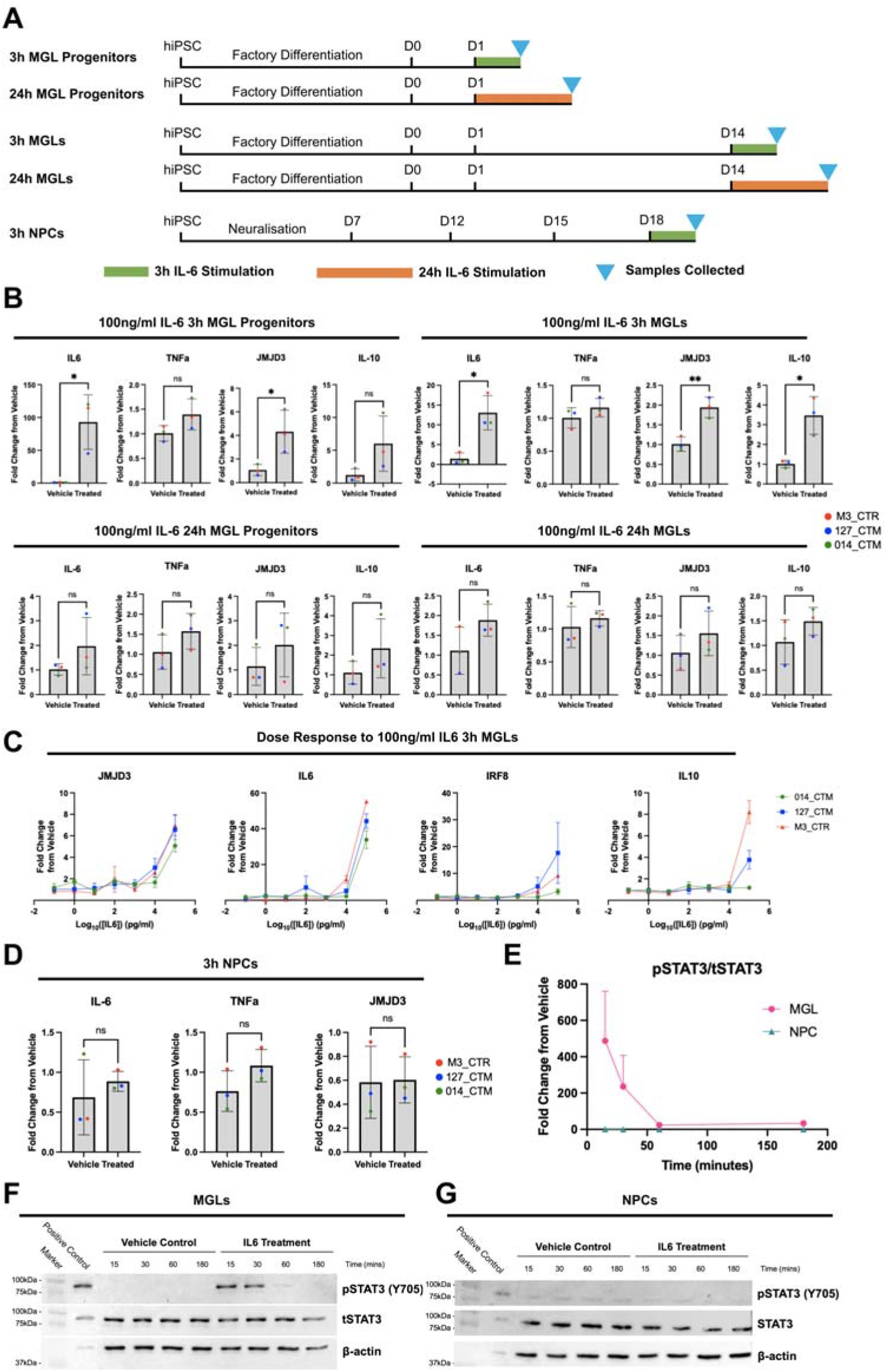
MGL monocultures respond to IL-6 in a dose and time dependent manner, NPC monocultures do not respond at all. (**A**) Schematic of MGL and NPC cell culture and RNA sample collection. (**B**) Transcripts of downstream IL-6 pathway genes *IL-6, TNFa, JMJD3* and *IL-10* were measured by qPCR in three male healthy control cell lines treated with 100ng/ml IL-6, averaged over three technical replicate clone cultures per donor unless stated otherwise, in the following conditions: MGL progenitors treated for 3h; MGLs treated for 3h; MGL progenitors treated for 24h, 014_CTM treated condition averaged from N=2 clones only; MGLs treated for 24h. Unpaired test results formatted as follows: *p < 0.05, **p < 0.01, ***p < 0.001, and ****p < 0.0001; not significant (ns). Bar graphs plotted as mean with standard deviation (SD) error bars, and points coloured by donor line: red (M3_CTR), blue (127_CTM) and green (014_CTM). (**C**) Dose response of MGLs to 7 doses of a 10-fold serial dilution of IL-6 from 100ng/ml to 0.1pg/ml. Three healthy male donors with n=3 harvest replicates per donor. *IL-10* 127_CTM_01 100pg/ml (Log_10_(2)) outlier removed and calculated form N=2 harvests. Fold change from vehicle calculated within line, but vehicle not plotted. (**D**) Transcripts of downstream IL-6 pathway genes *IL-6, TNFa* and *JMJD3* in NPCs treated for 3h, with unpaired t- test results formatted as before. *IL-10* transcripts were undetectable in NPC samples so data is not shown. (**E**) Quantification of pSTAT3/tSTAT3 signal in arbitrary (arb.) units from blots **F** and **G**, shown as a fold change ration from vehicle within cell type. (**F-G**) Immunoblotting for 88kDa pSTAT3/tSTAT3 in both vehicle and 100ng/ml IL-6 stimulated samples collected after 15, 30, 60 and 180mins, in MGL (**F**) and NPC (**G**) monocultures, with BV2 3h IL-6 100ng/ml treated positive control.

Having confirmed IL-6 triggers a transcriptional response associated with *cis*-IL6R signaling in MGLs, we next sought to determine the minimal concentration of IL-6 that would induce this response from day 14 MGLs in monoculture (**Figure 2C**). Of note, the mean concentration of IL-6 in maternal serum collected from second trimester mothers in a recent birth cohort study was reported to be 0.98±1.06 pg/ml, (Graham et al., 2018). At face value, our initial IL-6 concentration of 100 ng/ml IL-6 is therefore not representative of a physiologically relevant exposure based on the aforementioned human data (Graham et al., 2018). We exposed MGLs to several IL- 6 concentrations (range: 0.1 pg/ml to 100 ng/ml) and measured the expression of genes that were up regulated after 3h (*IL-6, IL-10* and *JMJD3*) by qPCR in MGLs. Complementing these data, we also measured expression of interferon regulatory factor 8 (*IFR8)*, a transcription factor known to regulate immune function and myeloid cell development (d’Errico et al., 2021; Tamura and Ozato, 2002). Expression of *IL-6, JMJD3, IL10* and *IRF8* were unaffected relative to vehicle control at all concentrations of IL-6 tested except 100 ng/ml, which elicited a clear increase in expression, which varied between donors as may be expected (**Figure 2C**, statistics in **Supplementary Table 15**). We therefore selected 100 ng/ml IL-6 for further experiments since this elicited a response in MGLs that can be measured at a single time-point. As already stated, this concentration is higher than would be observed *in vivo*, however it should be noted that in this study we are not developing a model system of MIA but investigating the response to a specific cytokine (IL-6) that plays a key role in MIA, as per our previous work on IFNγ (Bhat et al., 2022; Warre-Cornish et al., 2020).

Since forebrain NPCs, in contrast to MGLs, displayed a very low level of *IL6RA* expression, we sought to confirm whether forebrain NPCs in monoculture show any response to 100 ng/ml IL-6. At 3 hr post IL-6 exposure, forebrain NPCs did not significantly increase the expression of *IL-6, JMJD3* and *TNF*_α_ transcripts relative to vehicle controls (**Figure 2D**, statistics in **Supplementary Table 16**). Furthermore, the *IL-10* transcript was undetectable in all NPC samples, irrespective of treatment (*data not shown*). These data suggest that whilst IL-6 triggers *cis*-IL6R signaling in MGLs, this is not the case for forebrain NPCs at D18 *in vitro*.

To complement our gene expression analysis, we next assessed the time-scale of canonical STAT-3 signaling following IL-6 receptor stimulation at the protein level in both forebrain NPCs and MGLs. Formation of the IL-6/IL-6Ra/IL-6ST complex on the cell surface membrane results in phosphorylation of STAT3 by the protein kinase JAK (pSTAT3), which shuttles to the nucleus to enable subsequent transcription of STAT3 target genes (Wolf et al., 2014). We therefore collected protein samples at multiple time points following acute IL-6 exposure (100 ng/ml) of either NPCs or MGLs and performed western blotting for Y705-pSTAT3 and total STAT3 (**Figure 2E-G**). Quantification of the Y705-pSTAT3 ratio to total STAT3 (tSTAT3) indicated that IL-6 triggered a time-dependent increase in pSTAT3 relative to vehicle controls that peaked after 30mins in MGLs that was absent in D18 forebrain NPCs (**Figure 2E**, 2-way ANOVA: cell type F=8.564; p=0.0430, F=8.387; time p=0.0437, interaction F=8.363; p=0.0029).

### Acute IL-6 exposure elicits a transcriptional response in human microglia-like cells of relevance for schizophrenia

Our data thus far provide evidence suggesting that the canonical STAT-3 signaling pathway is activated in MGLs within 3hr of exposure to IL-6. To better characterize the transcriptional response of MGLs to this stimulus, we next performed bulk RNA-sequencing at 3 hr post IL-6 exposure (100 ng/ml) in MGLs generated from N=3 male neurotypical donor hiPSC lines. Principal component analysis (PCA) of the gene expression data reveals that samples clustered by treatment (**Figure 3A**), consistent with the heatmap clustering of the top 25 differentially expressed genes (DEGs) (**Figure 3B**). Overall, we found that 156 and 22 genes were up- and downregulated, respectively, following 3h IL-6 exposure (FDR < 0.05) (**Figure 3B-C**). Upregulated genes of note included *IRF8*, consistent with our qPCR data (**Figure 2C**) the NFkB subunit *REL*, heat shock proteins *HSPA1A/B* and the oxytocin receptor (*OXTR)*, although no effects of IL-6 were observed on key microglia markers *TMEM119, Iba1 and PU*.*1* (**Supplementary Figure 2**). The DESeq2 normalized counts for *IL-6Ra, IL-6ST* were also unaffected; confirming the expression of these receptors is independent of 3h IL-6 exposure (**Supplementary Figure 2**).

**Figure 3:**
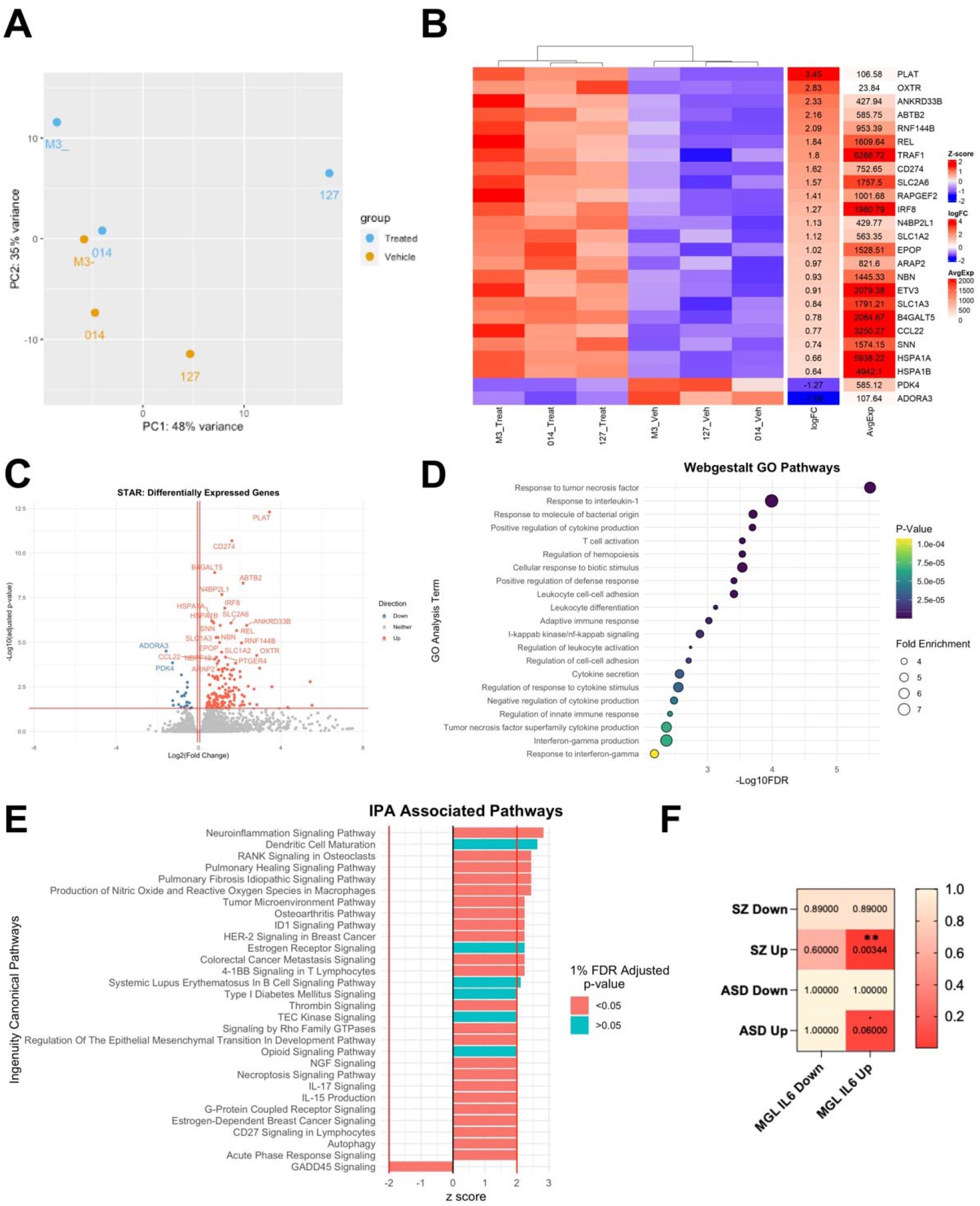
Acute IL-6 exposure elicits a transcriptional response in human microglia-like cells of relevance for schizophrenia. MGLs from 3 healthy male donors, pooled from 2 clone cultures each, were exposed to IL-6 or vehicle for 3h and collected for RNAseq. (**A**) PCA analysis of all 6 samples, coloured by vehicle (orange) or IL-6 treated (blue) condition and labelled by donor line: M3_CTR as M3-, 014_CTM as 014 and 127_CTM as 127. (**B**) Heatmap of top 25 most differentially expressed genes in the IL-6 3h MGL response, ranked by LogFC and clustering by treatment group. (**C**) Volcano plot of differentially expressed genes. Conditional axis set as follows: log2Foldchange > 0.06 and adjusted p-value < 0.05 coloured red; log2Foldchange < -0.06 and adjusted p-value < 0.05 coloured blue. The top 25 differentially expressed genes are labelled. (**D**) Webgestalt gene ontology analysis of upregulated 156 geneset only with an adjusted 1% FDR. GO terms ordered by –log10FDR, coloured by adjusted p-value and sized by the fold enrichment within each dataset. (**E**) IPA associated pathways, ranked by z-score and coloured by 1% FDR adjusted p-value. Only pathways with z-score > |2| are shown, with z-score > |2| conditional axes labelled in red. (**F**) Fisher’s exact test comparing genesets from ASC and SCZ post-mortem human patient tissue with up and down regulated genesets identified by RNAseq in this study (Cell Model). FDR plotted in heatmap with significant 5% FDR corrections formatted as follows: p < 0.1, *p < 0.05, **p < 0.01, ***p < 0.001, and ****p < 0.0001; not significant, not labelled.

Using only the 178 DEGs at 5% FDR, we carried out Webgestalt GO analyses splitting these into either upregulated (156) or downregulated genes (22). Across cellular components, biological processes, and molecular functions, 21 GO pathways were significantly associated with the 156-upregulated genes (1% FDR) (**Figure 3D**). These included the NFkB pathway (FDR=0.001), leukocyte differentiation (FDR<0.001) and cell-cell adhesion (FDR=0.002), response to cytokine stimuli such as IFNγ (FDR =0.007), production of IFNγ (q=0.004) and TNF superfamily (FDR =0.004) cytokines (**Figure 4D**). By contrast, no GO pathways were significantly associated with the 22 downregulated genes. Complementary GO analysis using the QIAGEN Ingenuity Pathway Analysis (IPA) software (Krämer et al., 2014) identified 30 associated pathways at a z-score threshold of > 2 to identify predicted activation or inhibition of a pathway, of which 24 passed 1% FDR correction (**Figure 4E**). The top activated pathways were neuroinflammation signaling (FDR <0.001), nitric oxide and reactive oxygen species (ROS) in macrophages (FDR <0.001), TNFRSF signaling in lymphocytes (4-1BB: FDR <0.001, CD27: FDR <0.001), epithelial-mesenchymal transition in development (FDR =0.001), G-protein coupled receptor signaling (FDR <0.001), IL-17 signaling (FDR =0.034) and a down regulation of GADD45 signaling (FDR =0.002). Overall, these complementary GO analyses provide evidence for a prototypical myeloid cell response after 3h of IL-6 stimulation, with NFkB pathway activation and downstream pathway changes to ROS, neuroinflammation, cell adhesion, cytokine secretion and TNFRSF signaling.

**Figure 4:**
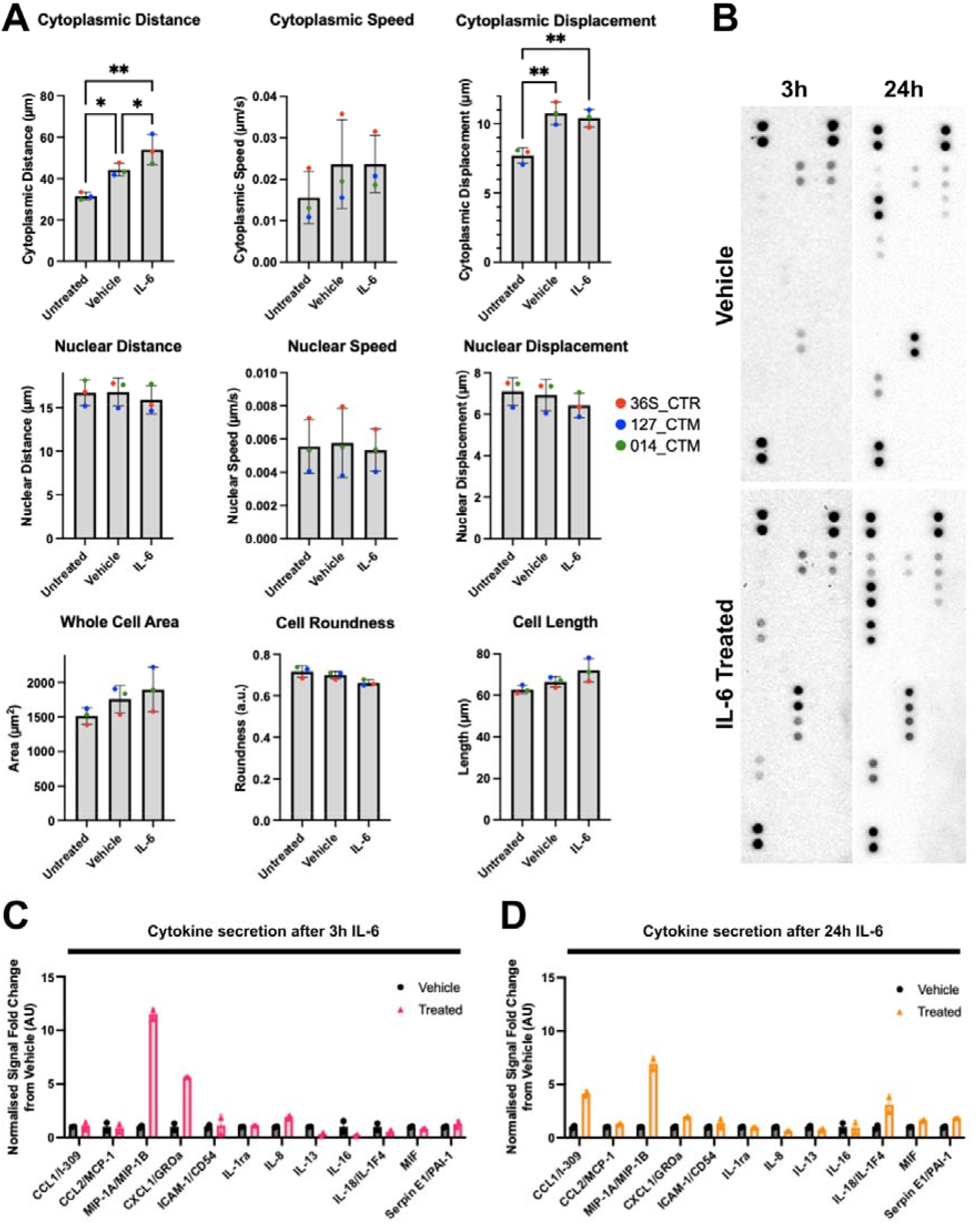
Acute IL-6 increases time-dependent changes in cytokine and chemokine secretion from human MGLs *in vitro*. MGL motility and secretome is altered in response to IL-6. (**A**) Metrics of MGL motility and morphology over 2h of live imaging, having been exposed to vehicle, IL-6 100ng/ml or untreated for 180 minutes. 5% FDR Benjamini method corrections formatted as follows: *p < 0.05, **p < 0.01, ***p < 0.001, and ****p < 0.0001; not significant not labelled. Bar graphs plotted as mean with standard deviation (SD) error bars, and points coloured by donor line: red (M3_CTR), blue (127_CTM) and green (014_CTM), all averaged from N=3 harvest replicates. (**B**) Representative images of dot blot cytokine profiles secreted from vehicle and IL-6 stimulated MGLs analysed using the human cytokine array. N=3 separate harvest media samples were pooled into one sample per condition from M3_CTR_37S. (**C-D**) Signal quantification of cytokine signals from dot blots presented in **B**, shortlisted for detectable cytokines and chemokines and split into 3h (**C**) and 24h (**D**) datasets. IL-6 not shown since it was artificially spiked by treatment. Each point represents a technical replicate of each signal point in arbitrary units, which were normalised to positive control reference spots and fold change calculated from averaged vehicle value within each time point.

We next investigated whether differentially expressed genes in our experimental conditions were enriched for genes differentially expressed in *post-mortem* brain samples originating from SZ or ASC cases (Gandal et al., 2018). We observed the 156 genes upregulated by IL-6 significantly overlapped with the genes upregulated in human *post-mortem* tissue from SZ cases (Gandal et al., 2018)(N genes upregulated in model = 156, N genes upregulated in cases = 2388, intersection size = 42 genes, P = 8.6e-04, FDR (corrected for four tests) = 0.00344, odds ratio = 1.8), but not with the genes downregulated in cases (P > 0.05). We found no overlap between the genes downregulated in our model with those up- and downregulated in cases (P > 0.05) (**Figure 3F**). We also observed a nominal enrichment between genes upregulated in the model with those upregulated in ASC cases (Gandal et al., 2018), but this did not pass multiple testing correction (**Figure 3F**) (N genes upregulated in model = 156, N genes upregulated in cases = 701, intersection size = 15 genes, P = 0.015, FDR (also corrected for four tests) = 0.06, odds ratio=2.0).

Finally, we investigated the overlap of up- and down-regulated genesets with microglia-specific module genesets using the MG Enrichment tool to compare data from published gene expression data in human microglia (Jao and Ciernia, 2021). When comparing our data to microglia gene sets derived from human tissue only, we found 31 modules were associated with the upregulated geneset from our cell model, and 12 with the down regulated geneset after 5% FDR correction. Notable modules that overlapped with our up and down-regulated genesets separately were the SCZ, ASD and Bipolar Disorder (BD) module (Up: intersection size = 44 genes, FDR = 7.10e-29, odds ratio = 13.5. Down: intersection size = 8 genes, FDR = 5.69e-06, odds ratio = 17.9) and the core human microglial signatures module (Up: intersection size = 26 genes, P = 3.29e-12, odds ratio = 6.3. Down: intersection size = 10 genes, FDR = 4.47e-08, odds ratio = 24.8). These data further establish the notion that IL-6- induced transcriptome of hiPSC-derived MGLs phenocopies not only core human microglia signatures, but is also of relevance for ASC, SCZ and BD disease states.

### Acute IL-6 exposure increases microglia motility and chemokine secretion in vitro

Both our RNAseq and qPCR data provide evidence for an increase in *IFR8* expression after IL-6 exposure in human MGLs. In mice, microglia-specific deletion of *IRF8* results in cells with fewer, shorter branches and reduced motility, consistent with the regulatory role of *IRF8* in microglia state (d’Errico et al., 2021). Furthermore, recent data from a mouse model of MIA provides evidence that IL-6 increases microglial motility *in vivo* (Ozaki et al., 2020). Based on these data we acquired live cell imaging data to record the effect of 3h exposure to IL-6 (100 ng/ml) on MGL whole cell (nuclear) motility, cytoplasmic specific (cytoplasm) motility and morphology, another known correlate of microglial function (Hanger et al., 2020). We observed that vehicle treatment was by itself sufficient to influence MGLs motility, as evidenced by an increase in cytoplasmic distance and displacement in both vehicle-and IL-6 treated cultures relative to untreated controls (**Figure 4A**, statistics in **Supplementary Table 17**). Critically however, IL-6 increased mean cytoplasmic distance relative to vehicle-treated controls, suggestive of increased cytoplasmic ruffling (**Figure 4A**, statistics in **Supplementary Table 17**). Nuclear motility, cell area and length were unchanged by either vehicle or IL-6 treatment (**Supplementary Figure 3**). We confirmed these effects were not due to differences in cell number or the movement of cells in or out of the field of view (statistics in **Supplementary Table 17**). Not only are these data consistent with increased *IRF8* expression (d’Errico et al., 2021) and our GO analysis, they may also suggest acute IL-6 exposure shifts MGLs into a state of enhanced environmental surveillance.

As several pathways related to cytokine secretion were specified during RNAseq GO analysis, we next aimed to gain an overview of cytokine secretion from IL-6 exposed MGLs by using a proteome profiler array, as previously described (Garcia-Reitboeck et al., 2018). Chemokine and cytokine release is clearly influenced by IL-6 stimulation at either 3 or 24h after exposure to IL-6 (100 ng/ml) (**Figure 4C-D** and **Supplementary Table 18**). Of the 36 cytokines and chemokines in the assay, 13 were above the limit of detection in culture supernatant. We excluded IL-6 since it was ectopically spiked into the media when the cells were stimulated with IL-6 (**Figure 4B**). Although semi-quantitative, changes in cytokine and chemokine secretion may be represented as fold changes from vehicle. MIP1A/B and CXCL1 were increased in supernatants from IL-6 exposed cultures at both time-points relative to vehicle-controls, although considerably less so after 24h; CCL1/2, Serpin-E1, MIF and IL-18 were increased only after 24h IL-6 stimulation with little difference observed at 3h; IL-8 presented higher secretion 3h after IL-6 simulation but not at 24h; the anti-inflammatory cytokines IL-13 and IL-16 were reduced at both time-points relative to vehicle-controls; and finally, IL-1ra, CCL2 and ICAM remained unchanged after IL-6 exposure at both time points. These alterations to the MGL secretome provide evidence that IL-6 stimulation of human MGLs leads to dynamic changes in specific inflammation-regulating chemokines and cytokines, that have time point specific secretion phases.

## Discussion

We characterized the cell-specific responses of MGL and NPC to acute IL-6 exposure using hiPSC lines obtained from male, neurotypical donors. Our data suggest two key findings; first, that hiPSC-derived MGL and NPC cells show clear differences in IL-6Ra cis-/trans-signaling capabilities and second, that exposure of MGLs to IL-6 recapitulates molecular and functional phenotypes of relevance for schizophrenia, consistent with evidence from genetic (Perry et al., 2021), blood biomarker (Allswede et al., 2020) and animal models (Smith et al., 2007) that connect this cytokine with increased risk for schizophrenia.

We observed that both hiPSC-derived forebrain NPCs and mature neurons express very low to undetectable levels of *IL-6Ra*, resulting in their inability to respond to IL-6 treatment in monoculture, as evidenced by the absence of STAT-3-phosphorylation and *IL-6, JMJD3* and *TNF*_α_ expression differences post-IL-6 exposure. These data are seemingly at odds with those from hiPSC-derived mature neurons derived using a similar differentiation protocol, in which IL-6 exposure resulted in transcriptional and morphological phenotypes (Kathuria et al., 2022b). One important difference between this work and our own is the age at which these cells were exposed to IL-6: specifically, D18 NPCs vs. D25-27 neurons (Kathuria et al., 2022b). Although we confirmed *IL-6Ra* expression is absent in forebrain neurons at D50, it may be possible that *IL-6Ra* mRNA is expressed transiently between days 25-50, or only protein levels are present, which is not detectable by RNAseq. Therefore, we cannot rule out that D25-27 neurons derived by this protocol can prompt an IL-6 response and that differences in the timing of IL-6 exposure may lead to differential results (Estes and McAllister, 2016). A second difference is the dose and length at which these cells were exposed to IL-6: 3h and 24h of 100ng/ml IL-6 herein as compared to 48h to 1μg/ml IL-6 (Kathuria et al., 2022b). It is important to note here that just as observed in animal MIA models, differences in the intensity and duration of immune activation will lead to variations in results. In this context we sought to find the minimum concentration of IL-6 that our cells would respond to *in vitro*, at least in monoculture, which is 100 ng/ml. Finally, we did not carry out RNAseq on our IL-6 exposed NPC RNA, so we cannot discount a non-canonical response to IL-6 that is independent of the IL-6Ra-STAT3 pathway at higher concentrations of IL-6. Overall, however, we do not see a response to IL-6 by forebrain NPCs under the conditions tested. Furthermore, our data strongly suggest microglial cells in co-culture with neural progenitors are necessary to study the influence of IL-6 on NPC-specific development when using D18 NPCs under the conditions tested in our study. In this context, we observed secretion of sIL-6Ra protein from MGLs but not NPCs, suggesting that in a co-culture paradigm, secretion of the sIL-6Ra by MGLs may enable NPCs to respond to IL-6 via *trans*-signaling *in vitro*. This is relevant since in the periphery, the cellular response to IL-6 is distinct depending on whether either *cis-* or *trans-*signaling pathways are activated (Su et al., 2017). In support of this view, data from a transgenic mouse model provide evidence that targeted inhibition of CNS *trans*-signaling via sIL6R mitigates a number of relevant neuropathological hallmarks previously associated with NDDs, including impaired neurogenesis, blood brain barrier leakage, vascular proliferation, astrogliosis and microgliosis (Campbell et al., 2014). These data support the view that IL6R *trans*-signaling is a relevant pathogenic mechanism of IL-6 in both glial and non-glial cell types (Campbell et al., 2014). Further studies are therefore now required to confirm in our system whether forebrain NPCs can activate trans-IL6 signaling in the presence of sIL6Ra protein.

Our second main observation is that the MGL response to acute IL-6 exposure *in vitro* phenocopies molecular data collected from cells and brain tissues of individuals with psychiatric disorders that have a putative neurodevelopmental origin, particularly schizophrenia (SZ). Specifically, we found the genes up regulated by hiPSC-derived MGLs after acute exposure to IL-6 significantly overlap with genes increased in *post-mortem* tissue from ASC, SZ and BD patients (Gandal et al., 2018). Key upregulated DEGs in this study, such as *HSPA1A/B, Rel* and *IRF8* are responsible for maintenance of microglial homeostasis, core microglial signatures and stress responses (Galatro et al., 2017; Gosselin et al., 2017; Olah et al., 2020), consistent with their differential expression after IL-6 stimulation. The observed overlap with SZ genesets however suggests links between IL-6 exposure, microglial signature pathways and SZ pathogenesis. For example, the NFκB pathway is known to be activated in both the periphery (Murphy et al., 2022) and in post-mortem tissue (Murphy et al., 2020) from SZ cases. Moreover, our observation that acute IL-6 stimulation is associated with up-regulation of the OXT receptor (OXTR) and associated signaling pathways, overlaps with evidence that polymorphisms in the *OXTR* gene are linked to the pathogenesis of both SZ (Broniarczyk-Czarniak et al., 2022; Nakata et al., 2021), but also ASC (de Oliveira Pereira Ribeiro et al., 2018; Francis et al., 2016). Intriguingly, studies in rodent primary microglia suggest OXT suppresses inflammatory responses following LPS stimulation *in vitro* (Inoue et al., 2019). Furthermore, in mice, treatment with an OXTR agonist reduces perinatal brain damage by specifically acting on microglia (Mairesse et al., 2019). Further studies are therefore required to investigate the role of OXTR signalling in regulating MGL responses to IL-6 in our human model system including studies in patient-derived cell lines.

Of note however, key microglial genes are also reported to be *down regulated* in both RNA seq and qPCR studies of human cortical *post-mortem* brain tissue from individuals with SZ (Gandal et al., 2018; Snijders et al., 2021). Taking an example directly relevant to our study, *IRF8* expression is reported to be *downregulated* in *post-mortem* human cortical brain tissue from individuals with SZ (Gandal et al., 2018; Snijders et al., 2021). In contrast to these data, we observed an increase of *IRF8* expression 3 hr acute IL-6 exposure, accompanied by increased cytoplasmic motility and time-dependent increases in chemokines and cytokines, findings consistent with the documented role of IRF8 in enabling microglia to adopt a pro-inflammatory gene signature in disease (Ransohoff and Engelhardt, 2012). This discrepancy could simply reflect differences related to methodology, since our human MGLs most closely resemble fetal microglia (Haenseler et al. 2017), whilst the *post-mortem* data are obviously influenced by ageing, duration of illness and potentially antipsychotic medication exposure, all of which may influence the results. Alternatively, it may reflect the fact that our MGLs were generated from hiPSC collected healthy donors and not individuals with SZ. Arguing against this however, Ormel and colleagues using PBMCs transdifferentiated to induced microglia (iM) identified two clusters of iM cells using mass cytometry, that were enriched only in donors with a diagnosis of SZ (Ormel at al. 2020). Of these, one cluster was characterized by *elevated* protein levels of CD68, Cx3cr1, HLA-DR, P2RY12, TGF- β1 and importantly, IRF-8 (Ormel et al., 2020). Further studies are therefore required to characterize how IL-6 impacts on specific microglia *states* of relevance to SZ via mass cytometry and/or single-cell sequencing approaches with appropriate functional assays.

Limitations of the current study should also be noted. As mentioned above, the response of each cell type presented here lies within the context of an acute IL-6 treatment in a monoculture, in the absence of a genetic background for schizophrenia or autism. In this context, birth cohort studies report the association between *average* IL-6 exposure across gestation, hence describing the impact of a cumulative exposure to IL-6 on brain and behaviour phenotypes (Graham et al., 2018). Furthermore, we have previously reported differential effects of acute IFN-gamma exposure on gene expression in forebrain NPCs from individuals with or without a diagnosis of SZ (Bhat et al., 2022). It will therefore be important to investigate both chronic IL-6 exposure and include patient-derived hiPSC lines in future studies to address these issues. In addition, our experiments were carried out using three individual clones per donor, from a total of N=3 donor lines, hence we cannot discount genotype specific IL-6 responses by the select few donors chosen here. Our sample size is however consistent with other studies of the impact of IL-6 on neurodevelopment using hiPSC models (Kathuria et al., 2022b). Combined with unique differentiation and/or cytokine exposure protocols reported by different laboratories, there is substantial risk that the reproducibility and hence, validity of mechanistic data from hiPSC models will be compromised (McNeill et al., 2020) Replication of our results by multiple groups using common hiPSC reference lines (e.g. corrected KOLF92) will therefore be an important advance for this field (Volpato and Webber, 2020).

In conclusion, hiPSC-derived MGLs can respond to IL-6 by either *cis-* or *trans-*signaling in monoculture, while NPCs in monoculture cannot due to the absence of *IL-6Ra* expression and sIL-6Ra secretion. The response of MGLs to IL-6 phenocopies molecular changes of relevance for SZ, consistent with the documented associations between IL-6 levels and risk for SZ (Perry et al., 2021). Our data also replicates key microglia findings from animal models of MIA in relation to motility and cytokine secretion, in a human model system. Collectively, our data underline the importance of studying microglial cells to understand the influence of IL-6 on human neurodevelopment and to elucidate cellular and molecular mechanisms that link early life immune activation to increased risk for psychiatric disorders with a putative neurodevelopmental origin.

## Materials and Methods

### Cell Culture

Participants were recruited and methods carried out in accordance with the ‘Patient iPSCs for Neurodevelopmental Disorders (PiNDs) study’ (REC No 13/LO/1218). Informed consent was obtained from all subjects for participation in the PiNDs study. Ethical approval for the PiNDs study was provided by the NHS Research Ethics Committee at the South London and Maudsley (SLaM) NHS R&D Office. HiPSCs were generated and characterized from a total of nine lines donated by three males with no history of neurodevelopmental of psychiatric disorders (**Supplementary Table 1**) as previously described (Adhya et al., 2021; Warre-Cornish et al., 2020) and grown in hypoxic conditions on Geltrex™ (Life Technologies; A1413302) coated 6-well NUNC™ plates in StemFlex medium (Gibco, A3349401) exchanged every 48 hours. For passaging, cells were washed with HBSS (Invitrogen; 14170146) and then passaged by incubation with Versene (Lonza; BE17-711E), diluted in fresh StemFlex and plated onto fresh Geltrex-coated 6-well NUNC™ plates. For specifics on cell culture differentiation, see supplementary. Both cell types were differentiated from hiPSCs using an embryonic MYB-independent method (Haenseler et al., 2017; Shi et al., 2012). Day 14 MGL monocultures for dose response experiments were exposed for 3 h to 100 ng/ml, 10 ng/ml, 1 ng/ml, 100 pg/ml, 10 pg/ml, 1 pg/ml, 0.1 pg/ml or 100 pM acetic acid vehicle and collected immediately for analysis. MGL progenitor (D1) and MGL (D14) cultures for single, high dose IL-6 stimulation received 100 ng/ml IL-6 (Gibco; PHC0066) or 100 pM acetic acid vehicle stimulation for either 3-or 24-hours and immediately collected for analysis. NPC cultures received 100 pM acetic acid vehicle or 100 ng/ml IL-6 (Gibco; PHC0066) on day 18 for 3h, then collected for analysis. Eighteen days after neural induction reflect early second trimester neurodevelopment, which corresponds to a known period of increased risk for offspring NDD in mothers with increased IL-6 serum concentrations (Estes and McAllister, 2016).

### RNA extraction, cDNA synthesis and quantitative PCR

Cells cultured for RNA extraction were collected in room temperature TRI Reagent™ Solution (Invitrogen; AM9738) and stored at -80ºC. RNA was extracted as directed per manufacturer’s instructions. Precipitation of RNA by 0.3 M Sodium-acetate and 100% ethanol at -80 ºC overnight was done to clean samples further, before resuspension in RNAse-free water. Nucleic acid content was measured using NanoDrop™ One. Reverse transcription of RNA to complementary DNA was carried out according to manufacturer’s instruction (SuperScript™ III Reverse Transcriptase Invitrogen 18080093 and 40 U RNaseOUT Invitrogen 10777019). qPCR was carried out using Forget-Me-Not™ EvaGreen® qPCR Master Mix (Biotium; 31041-1) in the QuantStudio 7 Flex Real-Time PCR System (Fisher), according to cycling parameters described in **Supplementary Table 6**. Cycle threshold (Ct) data were normalized to an average of *GADPH* and *RPL13* housekeeper expression Ct values.

### Western Blot

Cells were scraped on ice and collected in RIPA buffer (**Supplementary Table 7**), sonicated at 40% for 10 pulses, pelleted for 15 min at 4 ºC and proteins collected in supernatant. Protein concentration was quantified using the Pierce™ BCA protein assay kit (Thermo-Fisher; 23227). In preparation for SDS-PAGE separation, protein samples were denatured in Laemmli buffer and boiled at 95ºC for 5min. 2 μg of each protein sample was loaded into self-made 10% gels, alongside 5 μl of the Dual Color (BioRad #1610374) standards marker. Gels were run at 20mA for approximately 20 min, then increased to 100V until the samples reached the bottom of the unit (∼90min). Separated samples were transferred to a PVDF membrane and run overnight at 78mA in 4ºC. Blots were blocked in 5% BSA TBS-T for 1 hour at RT with agitation. Antibodies were diluted in blocking buffer; primary antibody incubation occurred overnight at 4ºC with agitation, and secondary antibody incubation at RT for 1 hour with agitation (**Supplementary Table 4**). Washes between antibody probes occurred in TBS-T at three 15min intervals. For visualization, ECL Western Blotting Substrate (GE Healthcare; RPN2106) was incubated on the blot at RT for 5min before image capture by the Bio-Rad Molecular Imager^®^ Gel Doc™ XR System.

### RNA Library Preparation and NovaSeq Sequencing

Total RNA extracted from 3h IL-6 treated day 14 MGLs was pooled from two clones per healthy male donor (n = 3 in total). The samples were submitted for sequencing at Genewiz Inc (South Plainfield, NJ). Libraries were prepared using a polyA selection method using the NEBNext Ultra II RNA Library Prep Kit for Illumina following manufacturer’s instructions (NEB, Ipswich, MA, USA) and quantified using Qubit 4.0 Fluorometer (Life Technologies, Carlsbad, CA, USA). RNA integrity was checked with RNA Kit on Agilent 5300 Fragment Analyzer (Agilent Technologies, Palo Alto, CA, USA). The sequencing libraries were multiplexed and loaded on the flowcell on the Illumina NovaSeq 6000 instrument according to manufacturer’s instructions. The samples were sequenced using a 2×150 Pair-End (PE) configuration v1.5. Image analysis and base calling were conducted by the NovaSeq Control Software v1.7 on the NovaSeq instrument. We obtained an average of 23.5 million 289-base pair paired-end reads per sample (**Supplementary Table 8**). All downstream analyses were carried out in R version 4.0.2 (R Core Team, 2020). FASTQ files were quality controlled using Fastqc (Wingett and Andrews, 2018) and subsequently aligned to the human reference genome (GRCh38) with STAR (Dobin et al., 2013). A count table was prepared and filtered for counts ≥ 1 using featureCounts (Liao et al., 2014) from the Rsubread (Yang Liao et al., 2019) package, version 2.4.3. Differential gene expression analysis was carried out using DESeq2 (Love et al., 2014) version 1.30.1 and the default Wald test. Subsequently, using the Benjamini-Hochberg (BH) method, only genes with adjusted P < 0.05 were considered differentially expressed and submitted for downstream analyses.

### Gene enrichments

Gene ontology (GO) analysis was carried out using WebGestalt (Yuxing Liao et al., 2019), where differentially expressed genes were tested for over representation of non-redundant cellular component, biological process and molecular function GO terms. This analysis used as a background list all genes considered expressed in our model, according to DESeq2s’s internal filtering criteria (i.e., adjusted P ≠ NA). Enrichment P-values were corrected for multiple testing using the BH method, and only terms with adjusted P < 0.05 were considered significant.

Outcomes from differential expression analysis were uploaded into the Qiagen Ingenuity Pathway Analysis (IPA) software (QIAGEN Inc., https://digitalinsights.qiagen.com/IPA) to identify canonical pathways. Analysis-ready genes were selected by p ≤ 0.05 and log-fold changes -0.06 ≤ or ≥ 0.06, resulting in 153 up-regulated and 22 down-regulated genes. Core analysis was filtered by human data and removed any cancer cell lines as reference from the IPA knowledge base (IPKB). Top 10 enriched canonical pathways were filtered by z-score ≥ |2|, an IPA measure of pathway directionality, and ordered by p-value adjusted by Benjamini-Hochberg corrections.

To calculate the overlap significance between genes up- or downregulated in our model with those up- or downregulated in *post-mortem* brain samples from schizophrenia or ASC cases (Gandal et al., 2018), we performed Fisher’s exact tests using the R package ‘GeneOverlap’ (Shen, 2021). We considered the number of genes expressed in our model and in the brain samples as genome size (n = 13,583). Multiple testing correction was applied using the FDR method, to correct overlap significance for the four tests performed for each disorder (upregulated in model vs. upregulated in cases, upregulated in model vs downregulated in cases, downregulated in model vs upregulated in cases, downregulated in model vs. downregulated in cases).

### Motility Assay

The motility of MGLs from donors M3_CTR_36S, 127_CTM_01 and 014_CTM_02, averaged from three harvests was measured by live imaging, with 6 technical repeat wells per condition. MGL progenitors were seeded onto a glass bottom 96 well plate (PerkinElmer) precoated with Poly-D-Lysine (Gibco; A3890401) at 22,000 cells/well and matured in MGL media for 14 days. On the day of imaging cells were exposed to 5 conditions: unstimulated, 100pM acetic acid vehicle for 30min, 100ng/ml IL-6 for 30min, 100pM acetic acid vehicle or 100ng/ml IL-6 for 3h. Prior to imaging, for the 3h treatment conditions a complete media change was done with microglia media containing either 100ng/ml IL-6 or 100pM acetic acid vehicle. Thirty min before imaging, complete media change was done on all remaining wells with microglia media containing either 100ng/ml IL-6, 100 pM acetic acid vehicle or neither (unstimulated). Simultaneously all cells were also stained for 30min with HCS NuclearMask™ Blue Stain (Invitrogen; H10325) and CellMask™ Orange Plasma membrane Stain (Invitrogen; C10045). Immediately before imaging, the media containing treatment and stain was removed and replaced with FluoroBrite™ DMEM (Gibco; A1896701) imaging media without phenol. Cells were imaged for 2h on an Opera Phoenix high throughput imaging system (Perkin Elmer) using a 20x objective over 5 consistent fields of view per well, and data was analyzed using Harmony High-Content Image analysis software (PerkinElmer).

### Media Cytokine Array

Day 14 MGL media samples collected after 3 and 24h of IL-6 exposure and pooled from one donor over 3 harvests were incubated with Proteome Profiler Human Cytokine antibody array membranes (R&D Systems; ARY005B), as per the manufacturers instructions. Dot blot signals were quantified using the Protein Array Analyzer Palette plug-in for ImageJ, and technical dot replicates averaged to one value. These values were backgrounded and normalized to positive reference controls on the dot blot.

### sIL-6R ELISA

The IL-6 Receptor (Soluble) Human ELISA Kit (Invitrogen; BMS214) was used as per the manufactures instructions to quantify soluble IL-6R expression in day 14 MGL and day 18 NPC vehicle/treated cell culture media. Optical density (OD) was blanked and measured at 450nm.

### Statistical Analysis

To account for variability between cultures, three distinct male donors considered as biological replicates, averaged from three technical replicate clone cultures per donor unless stated otherwise (**Supplementary Table 1**). The use of multiple clones per line enhanced reproducibility and ensured validation of results in each donor line. All statistical analyses were performed in Prism 9 for macOS version 9.3.1 (GraphPad Software LLC, California, USA), apart from RNAseq analyses which were carried out using the research computing facility at King’s College London, *Rosalind*, and R version 4.0.2 (R Core Team, 2020). Each specific test carried out is described in each corresponding figure legend, as well as the number of replicates hiPSC lines that make up each technical and biological replicates. Statistical summary tables can be found in the supplementary. When comparing means for more than 2 groups (**Supplementary Tables 11, 12 and 17**), one-way ANOVA was used. To test whether transcript expression changed after treating cells with IL-6, we performed unpaired t-tests (**Supplementary Tables 14 and 16**). When comparing means for two separate conditions (**Supplementary Tables 9, 10 and 15**), two-way ANOVA was used. *Post-hoc* testing was carried out using Benjamini method 5% or 1% FDR. P- and adjusted P-values <0.05 were considered as statistically significant. During GO analysis, 1% FDR cut off was chosen to concentrate the number of significantly associated pathways. During MGL quality control (**Supplementary Figure 1**), two-way ANOVAs comparing each gene expression with donor and time point demonstrated cell phenotype was not influenced by donor line (**Supplementary Tables 9 and 10**). Therefore, to reduce batch and reprogramming variability, biological replicates were considered as separate male donors which were averaged from N=3 technical replicates from either clone cultures of the same donor or separate MGL harvests, as described in figure legends. Statistics were not applied to media cytokine array data since the sample power was too low.

## Acknowledgements

The authors acknowledge use of the research computing facility at King’s College London, *Rosalind* (https://rosalind.kcl.ac.uk) and are thankful to George Chenell of the Wohl Cellular Imaging Centre at King’s College London for technical support during live imaging. ACMC, DPS and ACV acknowledge financial support for this study from the National Centre for the Replacement, Refinement and Reduction of Animals in Research (NC/S001506/1). The work (at King’s College, London) was also supported by the Medical Research Council (MRC) Centre grant (MR/N026063/1). AM, ACV and SK acknowledge support by the Neuro-Immune Interactions in Health & Disease Wellcome Trust PhD Training Programme (218452/Z/19/Z) at King’s College London.

## Notes

### Competing Interest Statement

The authors have declared no competing interest.

