## Supplementary Data for "Acute IL-6 exposure triggers canonical IL-6R signalling in hiPSC microglia, but not neural progenitor cells"

**Supplementary Methods and Results**

**Microglia-like cell culture and quality control**

MGL cell culture was differentiated as previously directed (Haenseler et al., 2017; van Wilgenburg et al., 2013). hiPSCs were collected by passaging, pelleted, and resuspended in StemFlex + 10µM Y-27632 to a final concentration of 4.0 x 10^6^ cells / ml, which was split evenly between wells in a 96-well low adherent plate. Over the following three days, the developing embryoid bodies (EB) underwent daily 75% medium exchanges with EB medium (**Supplementary Table 2**). EBs were then harvested wells, washed with HBSS and resuspended in factory medium (**Supplementary Table 2**). Approximately 75 EBs were added to one T75 flask and incubated in factory medium. Once a week, 5ml factory medium was added to the flask. After ~4 weeks, factories began to produce myeloid precursors. Cells were harvested weekly by removing supernatant alongside fresh factory medium replacements. Harvests of myeloid precursors were seeded on D0 for terminal differentiation to mature MGL cultures. Precursors were pelleted and seeded into a 6-well NUNC^TM^ plate at 1.5 x 10^6^ cells / well in Microglia medium (**Supplementary Table 2**) and adhered over the following 24 hours with 50% medium exchanges every 4 days until day 14. Quality control of MGLs at both protein and transcript level can be found in the supplementary results (**Supplementary Figure 1**).

*MYB-*independent myeloid precursors were successfully generated from a total of 9 lines; three healthy male donor hiPSC lines each with three clones replicate factories per donor (Haenseler et al., 2017; van Wilgenburg et al., 2013) (**Supplementary Table 1**). Mean precursor cell concentration per myeloid factory harvest was 327,000 cells/ml and mean viability was 68.6%, in line with data produced from different groups using the same protocols (Vaughan-Jackson et al., 2021) (**Supplementary Figure 1A**). Through differentiation, expression of microglial marker transcripts *P2RY12, MERTK, CX3CR1* and *TMEM119 increased,* and pluripotency transcript *Nanog* decreased, indicating an increasing microglial phenotype that matched previous hiPSC-derived MGL protocols (Haenseler et al., 2017) (**Supplementary Figure 1B, Supplementary Table 9**). Importantly, *CX3CR1* expression significantly increased from hiPSC expression after only 14 days of terminal differentiation, suggesting MGLs have an increased ability for neuron-glial signaling (Subbarayan et al., 2021)(**Supplementary Figure 1B**). *P2RY12* expression peaked at day 0 in the myeloid factory progenitor stage and decreased during terminal MGL differentiation (**Supplementary Figure 1B**). Expression of *MERTK* increased after differentiation from the hiPSC stage and remained consistently expressed through day 0, 1 and 14 of MGL differentiation. Finally, *TMEM119* expression remained constant through differentiation of hiPSC to MGL cells (**Supplementary Figure 1B**).

To ensure microglial-marker expression at a protein level and compare day 1 (MGL) progenitor and day 14 (MGL) phenotypes, cells were stained for PU.1 and TMEM119 (**Supplementary Figure 1C**). PU.1 is a key transcriptional factor that regulates microglial fate determination (Hoeffel and Ginhoux, 2015), whilst TMEM119 is identified by transcriptome studies as a robust microglial marker that effectively discriminates residential microglia from their closely related cell types, such as blood-derived macrophages in the human brain (Kaiser and Feng, 2019; Satoh et al., 2016). Importantly, the majority of cells (D1 81.4%, D14 85.2%) expressed both TMEM119 and PU.1 (**Supplementary Figure 1D**). This did not change between 1 and 14 days of terminal differentiation (PU1/TMEM119 expression p< 0.0001; Day p=0.9999; Interaction p=0.8869; Two-way ANOVA). Mean expression of PU.1, TMEM119 and cell area was not significantly different from 1 to 14 days of MGL differentiation from the myeloid factory, constantly across each healthy male donor (**Supplementary Figure 1E, Supplementary Table 10**).

In the MGL progenitor population at day 1 (n=338 cells), 81.4 ± 13.8% of cells expressed both TMEM119 and PU.1 and remained consistent through to mature MGL populations at day 14 (n=299), with 85.2 ± 9.8 % cells expressing both markers. Fewer cells expressed only PU.1 (day 1, 14.9 ± 13.9%; day 14, 10.8 ± 9.5%), and even fewer cells expressed neither markers (day 1, 3.7 ± 1.2 %; day 14, 3.9 ± 2.2%). No cells measured expressed only TMEM119.

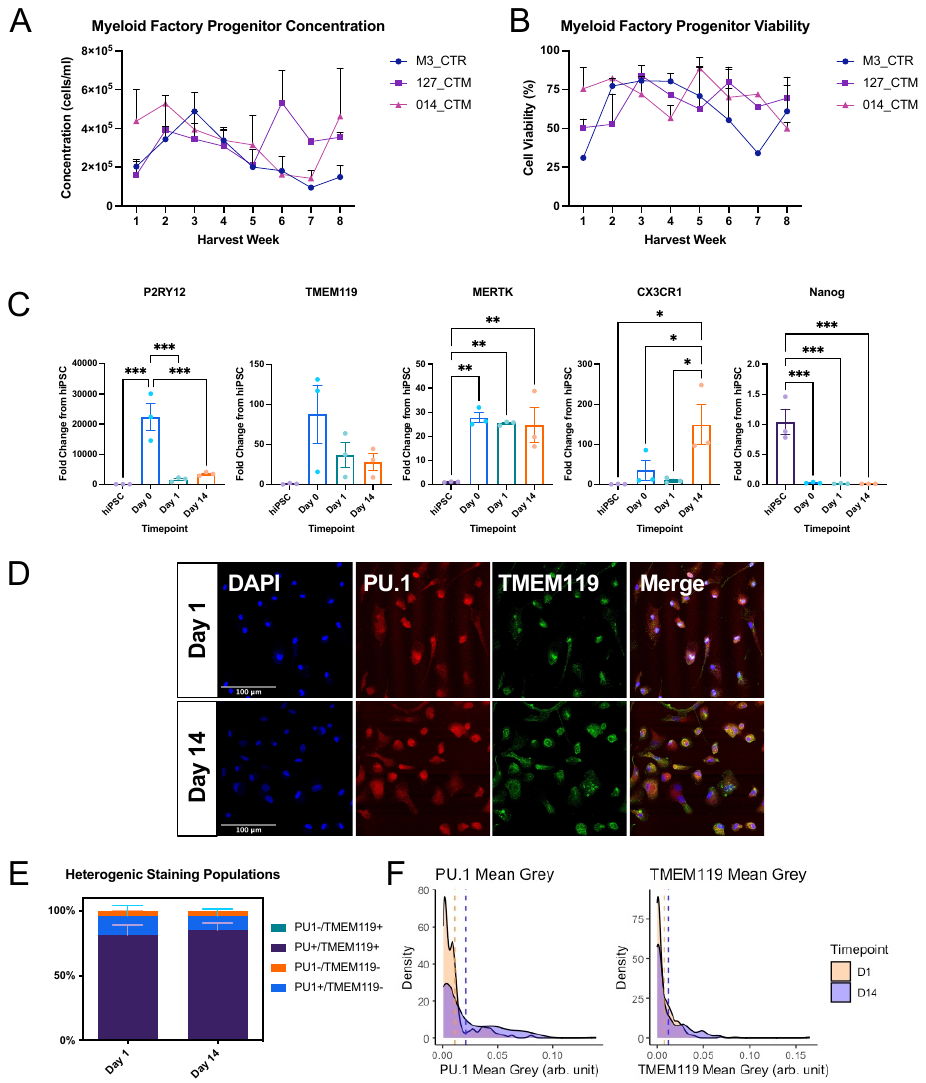

**Supplementary Figure 1: Expression of microglia markers in hiPSC derived MGLs.** Three male healthy control cell lines, averaged from three technical replicate clones per donor, unless stated otherwise. 5% FDR corrections formatted as follows: *p < 0.05, **p < 0.01, ***p < 0.001, and ****p < 0.0001; not significant not labelled. (**A**) Weekly harvest pre-MGL cell concentration from each donor line, N=3 clones per donor. (**B**) Weekly harvest pre-MGL cell viability from each donor line, N=3 clones per donor. **(C**) Differentiation time-course of microglial markers (*TMEM119, MERTK, CX3CR1*) and pluripotency marker (*Nanog*) by qPCR RNA samples at hiPSC, myeloid factory (day 0), MGL progenitor (day 1) and MGL (day 14) of differentiation. N=3 healthy male donors. 014_CTM Day 1 condition was averaged from N=2 clones only. (**D**) Representative confocal images of MGL progenitor (day 1) and MGL (day 14) collected by ICC, showing expression of both PU.1 (red) and TMEM119 (green). 127_CTM_01 line presented in this figure only. (**E**) Data extracted by CellProfiler from ICC PU.1/TMEM119 stained images showing proportions of PU.1/TMEM119 cell type populations; day 1 cell number=338, day 14 cell number=299, total cell number = 637. (**F**) Density plots distributing PU.1 and TMEM119 mean grey values. Vertical dashed line represents mean value for each timepoint.

**Neural progenitor cell culture**

Cultures of hiPSCs were neuralised using a modified dual SMAD inhibition protocol as previously defined to develop a mixture of excitatory and inhibitory forebrain neuronal subtypes (Shi et al., 2012). To commence neuralisation (day 0), hiPSC cultures were washed with HBSS and subsequently neuralised with induction medium (Supplementary Table 3). Cells continued to incubate with daily neuralisation medium changes. By day 7, a uniform neuroepithelial sheet appeared. These neuroepithelial cells were then collected and passaged to new plate. Hereafter day 8, SMAD inhibitors were omitted from culture medium and daily exchanges of maintenance medium proceeded (Supplementary Table 3). Neural maintenance medium was changed daily until day 18 of neuralisation. Neural passages were performed on day 12 and day 15 of neuralisation.

**Immunocytochemistry**

Cells were grown in 12-well plates on Geltrex-coated sterile coverslips and fixed with 4% formaldehyde (w/v; made in 4% sucrose PBS) for 20min at room temperature. Coverslips were washed twice in PBS, then simultaneously permeabilized and blocked with 2% normalized goat serum (NGS) PBS + 0.1 % Triton X-100 for 2 hours at RT. Antibodies were diluted in 2% NGS in PBS (**Supplementary Table 4**). Primary antibodies were incubated with cells overnight at 4ºC, and secondary antibodies were incubated with cells for 1 hour at RT, both in humidified chambers. Between incubations, coverslips were washed three times in PBS at 15min intervals. Finally, cells were incubated for 10min in PBS + DAPI (1:50,000). Coverslips were mounted onto glass slides with VECTASHIELD HardSet Antifade Mounting Medium (Vector Laboratories; H-1500-10). Confocal images were captured by a Leica SP5 laser scanning confocal microscope, with 405/488/594nm lasers and a 63x oil immersion lens (Leica, Wetzlar, Germany). Image analysis was performed in ImageJ and CellProfiler.

**cDNA Synthesis**

Reverse transcription of RNA to complementary DNA (cDNA) was carried out by incubating 1 µg RNA with 2.5µM Oligo(dT)_20_ and 0.5mM dNTP mix at 65 ºC for 5 min. Subsequently, 5X First-Strand Buffer, 5mM DTT, 200 U SuperScript^TM^ III Reverse Transcriptase (all Invitrogen; 18080093) and 40 U RNaseOUT (Invitrogen; 10777019) were added to the reaction mixture and incubated for 60 min at 50 ºC. The reaction was inactivated by a 15 min incubation at 70 ºC and diluted by a factor of 5 with RNAse-free water. Primers for intercalated dye (Forget-Me-Not™ EvaGreen® qPCR Master Mix; Biotium; 31041-1) quantitative PCR (qPCR) were designed using the IDT PrimerQuest^®^ Tool (**Supplementary Table 5**).

**Supplementary Table 1 –** Characteristics of 9 donor lines used throughout this study.

| Line Clone | Donor | Diagnosis | Reprogramming | Cohort | Age | Sex |
| --- | --- | --- | --- | --- | --- | --- |
| 014_CTM_02 | 014_CTM | Control | CytoTune™ Sendai | LEAP / StemBANCC | 18-30 | Male |
| 014_CTM_03 |  |  |  |  |  |  |
| 014_CTM_04 |  |  |  |  |  |  |
| M3_CTM_36S | M3_CTM | Control | CytoTune™ Sendai | EU-AIMS | 18-30 | Male |
| M3_CTM _37S |  |  |  |  |  |  |
| M3_CTM _38S |  |  |  |  |  |  |
| 127_CTM_01 | 127_CTM | Control | CytoTune™ Sendai | EU-AIMS | 50 - 60 | Male |
| 127_CTM_03 |  |  |  |  |  |  |
| 127_CTM_04 |  |  |  |  |  |  |

**Supplementary Table 2 –** Culture medias used during microglial cell type differentiation.

| Medium | Reagent | Final Concentration |
| --- | --- | --- |
| Embryoid Body (EB) | StemFlex (Gibco, A3349401) | 1X |
|  | BMP-4 | 50ng/ml |
|  | SCF | 20ng/ml |
|  | VEGF | 50ng/ml |
| Factory | X-VIVO 15 (Lonza; BE04-418F) | 1X |
|  | Glutamax (Life Technologies; 35050-038) | 2mM |
|  | IL-3 | 25ng/ml |
|  | M-CSF | 100ng/ml |
| Microglia | DMEM (Sigma; D6421) | 1X |
|  | N2 Supplement (Life Technologies; 17502-048) | 1% |
|  | Glutamax (Life Technologies; 35050-038) | 2mM |
|  | IL-34 | 100ng/ml |
|  | GM-CSF | 10ng/ml |

**Supplementary Table 3 –** Culture medias used during neural cell type differentiation.

| Medium | Reagent | Final Concentration |
| --- | --- | --- |
| N2 | DMEM (Sigma; D6421) | 1X |
|  | N2 Supplement (Life Technologies; 17502-048) | 1% |
|  | Glutamax (Life Technologies; 35050-038) | 2mM |
| B27 | Neurobasal Medium (Life Technologies; 21103-049) | 1X |
|  | B27 Supplement (Life Technologies; 17504-044) | 2% |
|  | Glutamax (Life Technologies; 35050-038) | 2 mM |
| Neuralisation | B27:N2 (as above) | 1:1 mixture |
|  | SB431542 (Cambridge Bioscience; ZRD-SB-50) | 10µM |
|  | LDN193189 (Sigma; SML0559) | 1µM |
| Maintenance | B27:N2 (as above) | 1:1 mixture |

**Supplementary Table 4 –** List of antibodies used during immunocytochemistry or western blotting.

| Epitope | Host | Dilution | Application | Manufacturer |
| --- | --- | --- | --- | --- |
| TMEM119 | Rabbit | 1:100 | ICC Primary | Abcam; ab185333 |
| PU.1 | Mouse | 1:100 |  | Santa Cruz; sc-390405 |
| Anti-Rabbit AlexaFluor 488 | Goat | 1:750 | ICC Secondary | Thermo Fischer Scientific; A11034 |
| Anti-Mouse AlexaFluor 568 | Goat | 1:750 |  | Thermo Fischer Scientific; A11031 |
| pSTAT3 (Y705) | Mouse | 1:1000 | WB Primary | Cell Signalling; 9138S |
| STAT3 | Rabbit | 1:1000 |  | Cell Signalling; 30835S |
| Beta-actin | Mouse | 1:5000 |  | Invitrogen; MA5-15739 |
| Anti-Rabbit HRP-conjugated | Goat | 1:5000 | WB Secondary | Thermo Fischer Scientific; 31460 |
| Anti-Mouse HRP-conjugated | Goat | 1:5000 |  | Thermo Fischer Scientific; 31430 |

**Supplementary Table 5 –** List of primers used during qPCR.

| Gene | Forward primer | Reverse primer |
| --- | --- | --- |
| Nanog | CCAACATCCTGAACCTCAGCTAC | GCCTTCTGCGTCACACCATT |
| TMEM119 | AGTCCTGTACGCCAAGGAAC | GCAGCAACAGAAGGATGAGG |
| MERTK | ACTTGCTGGTGGATGTTCC | ACTTGCTGGTGGATGTTCC |
| CX3CR1 | ACTTGCTGGTGGATGTTCC | ACTTGCTGGTGGATGTTCC |
| P2RY12 | GACAGGAGCTGCAGAACAGA | GTTGCCAAACCTCTTTGTGA |
| GP80 (IL-6R) | ACTTGCTGGTGGATGTTCC | ACTTGCTGGTGGATGTTCC |
| GP130 (IL-6ST) | GGCCTGAGTGAAACCCAAT | GGCCTGAGTGAAACCCAAT |
| IFNγR1 | GGTCTGTGAAGAGCCGTTGTC | GGTCTGTGAAGAGCCGTTGTC |
| IFNγR2 | GGTCTGTGAAGAGCCGTTGTC | GGTCTGTGAAGAGCCGTTGTC |
| TNFRSF1A | ACTTGCTGGTGGATGTTCC | ACTTGCTGGTGGATGTTCC |
| TNFRSF1B | GCCAGTGCGTTGGACAGAAG | CCACCAGGGGAAGAATCTGAG |
| IL17Ra | TGCGACTCCTGGACCAC | GTCAGGTTTCGAGGGTGAATC |
| TLR4 | CCCTGAGGCATTTAGGCAGCTA | AGGTAGAGAGGTGGCTTAGGCT |
| IL6 | GCGCTTGTGGAGAAGGAGT | TGGAGATGTCTGAGGCTCATT |
| JMJD3 | CCTTCTCACCTGTCCTGCTG | GGTCTTGGTGGAGAAGAGGC |
| IL10 | GTGGCGCTCCTGAGGTATGG | GTGGTACAGGTCCAAGGTCAC |
| TNFa | ACCAAGCCCGTGGTGAAG | TGACATCCTTGATGAAGAGCA |
| GADPH | GCAGCAACAGAAGGATGAGG | AACGTACTCAGCGCCAGCAT |
| RPL13 | CCCGTCCGGAACGTCTATAA | CCCGTCCGGAACGTCTATAA |

**Supplementary Table 6 –** Cycling parameters used during qPCR.

| Step | Temperature (ºC) | Time | Cycle |
| --- | --- | --- | --- |
| Initial denaturation | 95 | 10 mins | 1 |
| Denaturation | 95 | 15 sec | 40 |
| Annealing | 60 | 30 sec |  |
| Extension | 72 | 30 sec |  |

**Supplementary Table 7 –** RIPA buffer constitution, diluted in ddH_2_O.

| Reagent | Final Concentration |
| --- | --- |
| Tris pH 7.2 | 20mM |
| NaCl | 150mM |
| Triton X-100 | 1.0% |
| EDTA pH 8 | 5mM |
| SDS | 0.1% |
| Sodium Deoxycholate | 1% |
| AEBSF | 1mM |
| Pepstatin A | 1µg/ml |
| Leupeptin | 10µg/ml |
| Aprotinin | 10µg/ml |
| Ser/Thr phosphatase inhibitor cocktail | 1:100 |
| NaF | 25mM |

**Supplementary Table 8 –** Summary of read outputs per sample from RNAseq.

| Sample | M3-Veh | M3-Treat | 127-Veh | 127-Treat | 014-Veh | 014-Treat |
| --- | --- | --- | --- | --- | --- | --- |
| Unique read number | 20379588 | 21394231 | 24408798 | 29440355 | 25250460 | 21251473 |
| Average read length | 292 | 279 | 295 | 285 | 292 | 292 |

**Supplementary Table 9** – Two-way ANOVA of qPCR microglial marker gene expression with N=3 donors, each with N=3 clones.

| Gene | Source of Variation | DF | F (DFn, DFd) | P value | P value summary |
| --- | --- | --- | --- | --- | --- |
| P2RY12 | Interaction | 6 | F (6, 23) = 0.6872 | 0.6620 | ns |
|  | Timepoint | 3 | F (3, 23) = 5.335 | 0.0061 | ** |
|  | Donor | 2 | F (2, 23) = 0.9417 | 0.4045 | ns |
| TMEM119 | Interaction | 6 | F (6, 23) = 0.4903 | 0.8089 | ns |
|  | Timepoint | 3 | F (3, 23) = 1.112 | 0.3647 | ns |
|  | Donor | 2 | F (2, 23) = 2.905 | 0.0750 | ns |
| MERTK | Interaction | 6 | F (6, 23) = 1.302 | 0.2957 | ns |
|  | Timepoint | 3 | F (3, 23) = 11.49 | <0.0001 | **** |
|  | Donor | 2 | F (2, 23) = 4.167 | 0.0285 | * |
| CX3CR1 | Interaction | 6 | F (6, 23) = 0.2142 | 0.9685 | ns |
|  | Timepoint | 3 | F (3, 23) = 3.398 | 0.0349 | * |
|  | Donor | 2 | F (2, 23) = 0.5630 | 0.5771 | ns |
| Nanog | Interaction | 6 | F (6, 23) = 0.06092 | 0.9989 | ns |
|  | Timepoint | 3 | F (3, 23) = 33.38 | <0.0001 | **** |
|  | Donor | 2 | F (2, 23) = 0.03520 | 0.9655 | ns |

**Supplementary Table 10** – Two-way ANOVA of ICC microglial marker TMEM119 and PU.1 protein expression with N=3 donors, each with N=3 clones.

| Measurement | Source of Variation | DF | F (DFn, DFd) | P value | P value summary |
| --- | --- | --- | --- | --- | --- |
| PU.1 Mean Grey | Interaction | 2 | F (2, 12) = 0.8557 | P=0.4494 | ns |
|  | Time Factor | 1 | F (1, 12) = 1.506 | P=0.2432 | ns |
|  | Donor Factor | 2 | F (2, 12) = 0.6844 | P=0.5230 | ns |
| TMEM119 Mean Grey | Interaction | 2 | F (2, 12) = 0.8489 | P=0.4520 | ns |
|  | Time Factor | 1 | F (1, 12) = 1.304 | P=0.2757 | ns |
|  | Donor Factor | 2 | F (2, 12) = 1.344 | P=0.2974 | ns |
| Mean Whole Cell Area | Interaction | 2 | F (2, 12) = 0.3446 | P=0.7153 | ns |
|  | Time Factor | 1 | F (1, 12) = 0.9985 | P=0.3374 | ns |
|  | Donor Factor | 2 | F (2, 12) = 0.1248 | P=0.8838 | ns |

**Supplementary Table 11** – One-way ANOVA of qPCR cytokine receptor expression in MGLs with N=3 donors, averaged to one point from three clones.

| Gene | DF | F (DFn, DFd) | P value | P value summary |
| --- | --- | --- | --- | --- |
| INFyR1 | 3 | F (3, 8) = 48.40 | P<0.0001 | **** |
| IFNyR2 | 3 | F (3, 8) = 22.12 | P=0.0003 | *** |
| TNFRSF1A | 3 | F (3, 8) = 12.41 | P=0.0022 | ** |
| TNFRSF1B | 3 | F (3, 8) = 3.571 | P=0.0666 | ns |
| IL-6a (GP80) | 3 | F (3, 8) = 10.95 | P=0.0033 | ** |
| IL-6ST (GP130) | 3 | F (3, 8) = 10.06 | P=0.0043 | ** |
| IL-17Ra | 3 | F (3, 8) = 3.008 | P=0.0946 | * |

**Supplementary Table 12** – One-way ANOVA of qPCR cytokine receptor expression in NPCs with N=3 donors, averaged to one point from three clones.

| Gene | DF | F (DFn, DFd) | P value | P value summary |
| --- | --- | --- | --- | --- |
| INFyR1 | 3 | F (3, 8) = 0.4339 | P=0.7346 | ns |
| IFNyR2 | 3 | F (3, 8) =0.6834 | P=0.5867 | ns |
| TNFRSF1A | 3 | F (3, 8) = 1.325 | P=0.3324 | ns |
| TNFRSF1B | 3 | F (3, 8) = 6.972 | P=0.0127 | * |
| IL-6a (GP80) | 3 | F (3, 8) = 3.008 | P=0.0946 | ns |
| IL-6ST (GP130) | 3 | F (3, 8) = 2.464 | P=0.1369 | ns |
| IL-17Ra | 3 | F (3, 8) = 0.2049 | P=0.8902 | ns |

**Supplementary Table 13** – Validation of *IL-6Ra (gp80)* and *IL-6ST (gp130)* expression in previous transcriptomic datasets.

| Paper | Species | Age | Method | Sample | GP80 | GP130 |
| --- | --- | --- | --- | --- | --- | --- |
| Zhang et al., 2014 | Human | Fetal/  adult | RNAseq | Microglia/ macrophage | 8.12 FPKM | 80.06 FPKM |
|  |  |  |  | Neurons | 0.28 FPKM | 18.59 FPKM |
|  |  |  |  | Fetal Astrocytes | 0.65 FPKM | 9.2 FPKM |
| Miller et al., 2014 | Human | Midfetal | Micro-  array | Bulk | 0.41 RPKM | 5.35 RPKM |

**Supplementary Table 14** – Unpaired t-test p-value statistic of IL6, IL-10, JMJD3 and TNFa expression changes in MGL progenitor and MGL IL-6 treated samples over 3 and 24h.

| Cell Type | Gene | 3h P value | 3h P value Summary | 24h P value | 24h P value Summary |
| --- | --- | --- | --- | --- | --- |
| MGL Progenitor | IL-6 | 0.0186 | * | 0.2391 | ns |
|  | TNFa | 0.1312 | ns | 0.2202 | ns |
|  | JMJD3 | 0.0401 | * | 0.3748 | ns |
|  | IL-10 | 0.1264 | ns | 0.2529 | ns |
| MGL | IL-6 | 0.0118 | * | 0.1378 | ns |
|  | TNFa | 0.2620 | ns | 0.5259 | ns |
|  | JMJD3 | 0.0073 | ** | 0.3004 | ns |
|  | IL-10 | 0.0120 | * | 0.2419 | ns |

**Supplementary Table 15** – Two-way ANOVA of qPCR dose response gene fold changes from vehicle, with N=3 donors averaged to one point from three clones, over 5 doses.

| Gene | Source of Variation | DF | F (DFn, DFd) | P value | P value summary |
| --- | --- | --- | --- | --- | --- |
| IL-6 | Interaction | 12 | F (12, 42) = 4.782 | P<0.0001 | **** |
|  | Dose Factor | 6 | F (6, 42) = 168.9 | P<0.0001 | **** |
|  | Donor Factor | 2 | F (2, 42) = 4.685 | P=0.0146 | * |
| IL-10 | Interaction | 12 | F (12, 41) = 13.21 | P<0.0001 | **** |
|  | Dose Factor | 6 | F (6, 41) = 35.11 | P<0.0001 | **** |
|  | Donor Factor | 2 | F (2, 41) = 11.48 | P=0.0001 | *** |
| JMJD3 | Interaction | 12 | F (12, 42) = 0.9209 | P=0.5352 | ns |
|  | Dose Factor | 6 | F (6, 42) = 32.98 | P<0.0001 | **** |
|  | Donor Factor | 2 | F (2, 42) = 0.4621 | P=0.6331 | ns |
| IRF8 | Interaction | 12 | F (12, 42) = 1.083 | P=0.3982 | ns |
|  | Dose Factor | 6 | F (6, 42) = 4.748 | P=0.0009 | *** |
|  | Donor Factor | 2 | F (2, 42) = 1.666 | P=0.2013 | ns |

**Supplementary Table 16** – Unpaired t-test p-value statistic of IL6, JMJD3 and TNFa expression changes in NPC IL-6 treated samples over 3h.

| Gene | 3h P Value | P value summry |
| --- | --- | --- |
| IL-6 | 0.5161 | ns |
| TNFa | 0.1634 | ns |
| JMJD3 | 0.9275 | ns |

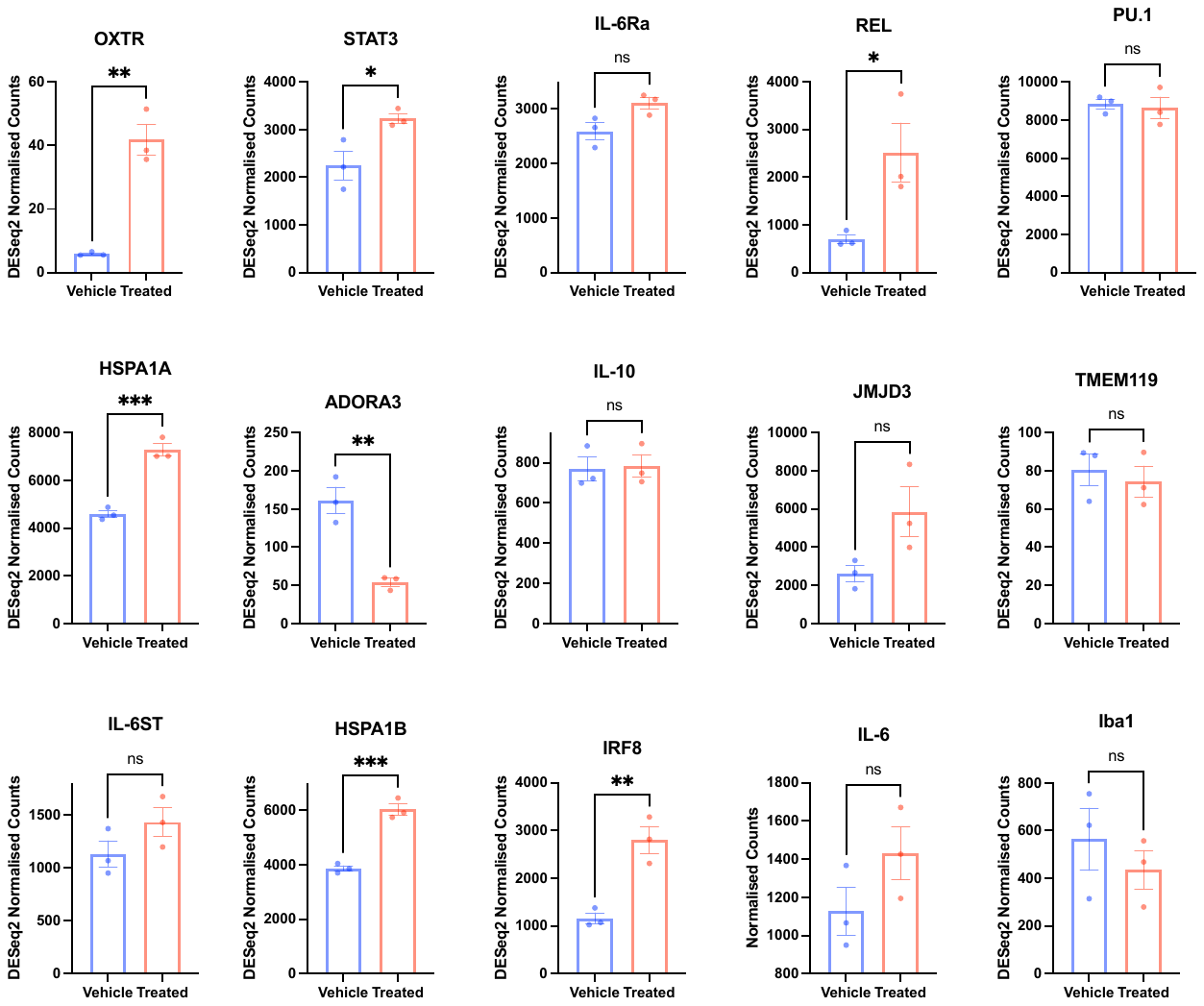

**Supplementary Figure 2 –** DESeq2 normalized counts of genes of interest after bulk RNAseq analysis. Bar graphs plotted as mean with standard error of the mean (SEM) error bars, with N=3 donor replicates. Unpaired test results formatted as follows: *p < 0.05, **p < 0.01, ***p < 0.001, and ****p < 0.0001; not significant (ns).

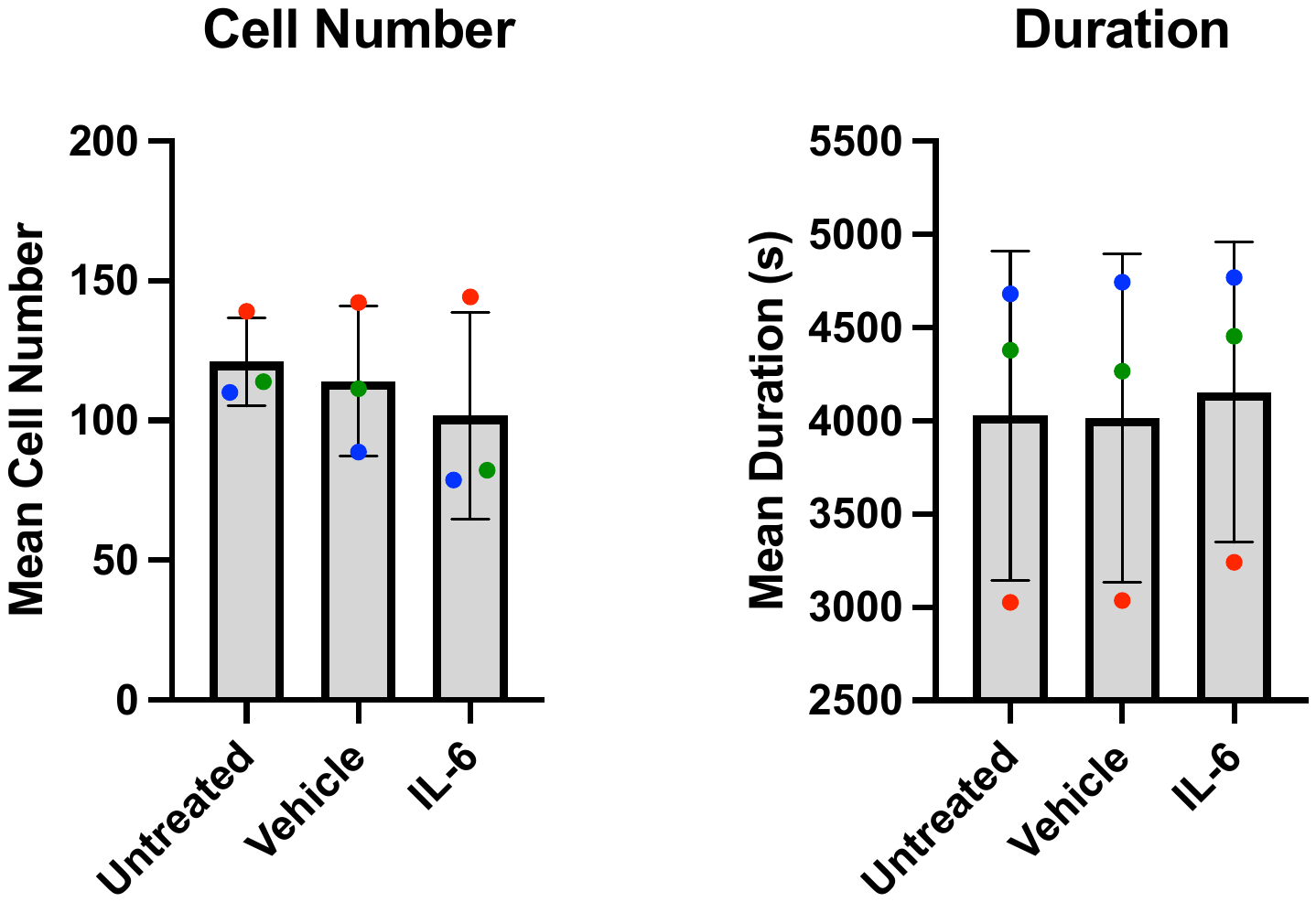

**Supplementary Figure 3 –** Motility assay quality control showing mean cell number per field of view and the mean duration of cells was unchanged by each condition. Bar graphs plotted as mean with standard deviation (SD) error bars, and points coloured by donor line: red (M3_CTR), blue (127_CTM) and green (014_CTM), all averaged from N=3 harvest replicates.

**Supplementary Table 17** – One-way ANOVA of Motility Assay conditions, with N=3 male donors averaged each from N=3 separate MGL harvest technical replicates.

| Measurement | F | P value | P value Summary |
| --- | --- | --- | --- |
| Cytoplasm Distance | 17.35 | 0.0032 | ** |
| Cytoplasm Speed | 0.9725 | 0.4307 | ns |
| Cytoplasm Displacement | 18.24 | 0.0028 | ** |
| Nuclear Distance | 0.3008 | 0.7508 | ns |
| Nuclear Speed | 0.04809 | 0.9534 | ns |
| Nuclear Displacement | 0.8013 | 0.4915 | ns |
| Area | 2.191 | 0.1930 | ns |
| Roundness | 5.425 | 0.0451 | * |
| Length | 4.632 | 0.0607 | ns |
| Number | 0.3693 | 0.7059 | ns |
| Duration | 0.02407 | 0.9763 | ns |

**Supplementary Table 18** – Mean signal values from cytokine profiler dot blots, backgrounded and normalised to positive control reference dots.

| Cytokine | 3h Vehicle | 3h Treated | 24h Vehicle | 24h Treated |
| --- | --- | --- | --- | --- |
| CCL1/I-309 | 0.01 | 0.01 | 0.05 | 0.22 |
| CCL2/MCP-1 | 0.02 | 0.02 | 0.75 | 0.98 |
| MIP-1A/MIP-1B | 0.01 | 0.15 | 0.09 | 0.65 |
| CCL5/RANTES | 0.01 | 0.00 | 0.00 | 0.00 |
| TNFSF5 | 0.01 | 0.00 | 0.00 | 0.00 |
| C5a | 0.00 | 0.00 | 0.00 | -0.01 |
| CXCL1/GROa | 0.01 | 0.08 | 0.22 | 0.44 |
| CXCL10/IP-10 | 0.01 | 0.00 | 0.01 | 0.00 |
| CXCL11/I-TAC | 0.00 | 0.00 | 0.01 | 0.00 |
| CXCL12/SDF-1 | 0.00 | 0.00 | 0.00 | 0.00 |
| G-CSF | 0.00 | 0.00 | 0.00 | 0.00 |
| GM-CSF | 0.01 | 0.01 | 0.00 | 0.00 |
| ICAM-1/CD54 | 0.01 | 0.01 | 0.01 | 0.01 |
| IFN-y | 0.00 | 0.00 | 0.01 | 0.00 |
| IL-1a | 0.00 | 0.00 | 0.00 | 0.00 |
| IL-1b | 0.00 | 0.00 | 0.00 | 0.00 |
| IL-1ra | 0.19 | 0.22 | 0.09 | 0.09 |
| IL-2 | 0.00 | 0.00 | 0.00 | 0.00 |
| IL-4 | 0.01 | 0.00 | 0.00 | 0.00 |
| IL-5 | 0.01 | 0.00 | 0.00 | -0.01 |
| IL-6 * | 0.00 | 0.53 | 0.01 | 0.55 |
| IL-8 | 0.11 | 0.21 | 0.73 | 0.47 |
| IL-10 | 0.00 | 0.00 | 0.00 | 0.00 |
| IL-12 p70 | 0.00 | 0.00 | 0.00 | 0.00 |
| IL-13 | 0.01 | 0.00 | 0.01 | 0.00 |
| IL-16 | 0.01 | 0.00 | 0.00 | 0.00 |
| IL-17A | 0.01 | -0.01 | 0.00 | 0.00 |
| IL-17E | 0.01 | 0.00 | 0.00 | 0.01 |
| IL-18/IL-1F4 | 0.01 | 0.01 | 0.01 | 0.02 |
| IL-21 | 0.00 | 0.00 | 0.01 | 0.01 |
| IL-27 | 0.00 | 0.00 | 0.00 | 0.01 |
| IL-32a | 0.00 | 0.00 | 0.00 | 0.01 |
| MIF | 0.19 | 0.16 | 0.11 | 0.19 |
| Serpin E1/PAI-1 | 0.02 | 0.02 | 0.07 | 0.13 |
| TNFa | 0.00 | 0.00 | 0.00 | 0.01 |
| TREM-1 | 0.00 | 0.00 | 0.00 | 0.01 |
